# NFIB and TCF4 converge to regulate the expression of genes involved in neuronal differentiation during neocortical development

**DOI:** 10.64898/2026.11.10.765673

**Authors:** Jonathan W.C. Lim, Emran Hossen, Nathan A. Mundell, Jesse E. Antoniak, Suranjana Pal, Henry Burgess, Jens Bunt, Linda J. Richards

## Abstract

*NFIB* or *TCF4* haploinsufficiency results in distinct neurodevelopmental disorders that share several characteristic overlapping features. Recent studies suggest that these transcription factors converge by binding to the same cis-regulatory elements and thereby regulate a common set of target genes. Here, we sought to determine whether in mice, NFIB and TCF4 converge during neocortical development. NFIB and TCF4 are co-expressed in radial glial cells and postmitotic neurons. Transcriptomic analyses of *Nfib* and *Tcf4* knockout mice uncovered an overlapping set of downregulated genes involved in neuronal differentiation. Analyses of genome-wide binding further demonstrate that NFIB and TCF4 largely bind to the same regulatory elements. In luciferase assays, NFIB and TCF4 elicit distinct effects on transcriptional activity that differ between the regulatory elements and cell types that we tested. We propose that this heterogeneity is a means through which NFIB and TCF4 function cooperatively to fine-tune target gene expression. Our study provides critical evidence for the molecular convergence of NFIB and TCF4. This indicates that the human syndromes resulting from mutations in these genes may have overlapping underlying etiologies.

**Summary statement:** NFIB and TCF4 regulate similar target genes during neocortical development by binding to the same regulatory elements. This convergence could contribute to common neurological phenotypes in *NFIB* and *TCF4* haploinsufficiency.

## Introduction

Neurodevelopmental disorders (NDDs) are a broad and complex class of disorders with a strong underlying genetic component. Next-generation sequencing has brought about the rapid discovery of many genes linked to NDDs (Bowling et al., 2017; Wright et al., 2023; Wright et al., 2015). However, functional studies aimed at investigating the underlying mechanistic etiology have struggled to keep pace, as such studies demand significantly more resources. An appealing approach to overcoming this challenge is to identify causal mechanisms linked to functional pathways where multiple genes may molecularly converge. One such functional pathway found to be implicated in NDDs is transcriptional regulation (Hormozdiari et al., 2015; Parenti et al., 2020) Haploinsufficiency of transcriptional regulators is a common cause of NDDs (Zug, 2022). Further, the cis-regulatory elements that transcription factors bind to often contain multiple copies of many transcription factor binding sites. Therefore, NDD-associated transcription factors could functionally converge by binding to the same cis-regulatory elements to regulate a common set of downstream genes. Identifying such transcription factors may be a means for revealing molecular mechanisms responsible for phenotypes shared between multiple NDDs.

We previously described a novel NDD caused by *NFIB* haploinsufficiency (Schanze et al., 2018). This disorder is characterized by developmental delay, mild-to-moderate intellectual disability, macrocephaly, and occasional corpus callosum dysgenesis (CCD). Affected individuals also display mild dysmorphic facial features that include a long face with high forehead, down-slanting palpebral fissures, a narrow nasal bridge, anteverted nares, and a long and smooth philtrum. Studies aimed at understanding the underlying etiology of neurological phenotypes are currently lacking. In mice, NFIB is first expressed in radial glial cells and differentiating neurons at E12.5 (Plachez et al., 2008), but downstream targets of NFIB in these cell types are predominantly unknown (Betancourt et al., 2014; Bunt et al., 2017; Piper et al., 2014; Steele-Perkins et al., 2005). Studies focused on chromatin modifiers (Martins-Costa et al., 2024; Quevedo et al., 2019) and interneuron development (Bright et al., 2025) suggests potential molecular convergence between NFIB and the basic helix-loop-helix transcription factor TCF4.

Whether NFIB and TCF4 molecularly converge is intriguing, as the phenotypes associated with haploinsufficiency of either gene are predominantly distinct. *TCF4* haploinsufficiency causes Pitt-Hopkins syndrome, which is associated with developmental delay, severe intellectual disability, microcephaly, and occasional CCD (Amiel et al., 2007; Whalen et al., 2012; Zollino et al., 2019; Zweier et al., 2007). Affected individuals also display facial features that are noticeably different from *NFIB* haploinsufficiency. These include a narrow forehead, a wide nasal bridge and bulbous nasal tip, full cheeks, and a wide mouth and full lips. A particularly appealing hypothesis is that similar phenotypes such as developmental delay, intellectual disability, and CCD may arise due to shared downstream targets, whereas phenotypes observed in only one disorder may be mediated by unique targets. Here, we examined mouse models to determine whether NFIB and TCF4 functionally and molecularly converge by regulating similar target genes during neocortical development. We find evidence for this such as a significant overlap in misregulated genes in knockout mice and almost identical genome-wide binding. Luciferase reporter assays indicate that when co-expressed, NFIB and TCF4 may bring about complex combinatorial outcomes that are distinct between different regulatory elements and in different cell types. This may be a means through which NFIB and TCF4 elicit pleiotropic effects despite overall binding to the same target genes.

## Results

### NFIB and TCF4 are co-expressed in radial glia and postmitotic neurons during neocortical development

NFIB and TCF4 expression have been examined in separate studies but their co-expression is unknown (Betancourt et al., 2014; Bunt et al., 2015; Bunt et al., 2017; Burette et al., 2024; Jung et al., 2018; Li et al., 2019; Mesman et al., 2020; Plachez et al., 2008). We performed immunofluorescence staining of wildtype coronal sections using antibodies specific for NFIB and TCF4, as well as CTIP2 or SOX2 to co-label early-born neurons or radial glial cells, respectively. NFIB expression is detected from E12.5 (Betancourt et al., 2014; Plachez et al., 2008), with lower expression within the ventricular zone and higher expression in postmitotic neurons occupying the nascent cortical plate (Fig. 1A). TCF4 is lowly expressed at E12.5 as reported previously (Fig. 1A) (Mesman et al., 2020). At E13.5, TCF4 expression is generally comparable to NFIB. Both proteins are co-expressed at lower levels in SOX2-positive radial glial cells and at higher levels in postmitotic neurons, particularly those within the cortical plate (Fig. S1 and yellow arrows in Fig. 1B’, respectively). From E14.5 onwards, while some cells highly co-express both NFIB and TCF4, other cells predominantly expressed one or the other. NFIB continues to be highly expressed within the cortical plate (Fig. 1C,D; white arrows in Fig. 1C’), but at E17.5, is only highly expressed within the deeper layers (Fig. 1E). On the other hand, TCF4 is most highly expressed within the intermediate zone at E14.5 and E15.5 (Fig. 1C,D; yellow arrowheads in Fig. 1C’) and then in the upper cortical layers at E17.5 (Fig. 1E). This suggests that NFIB and TCF4 expression arise at neurogenesis and then each contribute more specifically to the generation of the deeper and upper layers of the neocortex, respectively.

**Figure 1.**
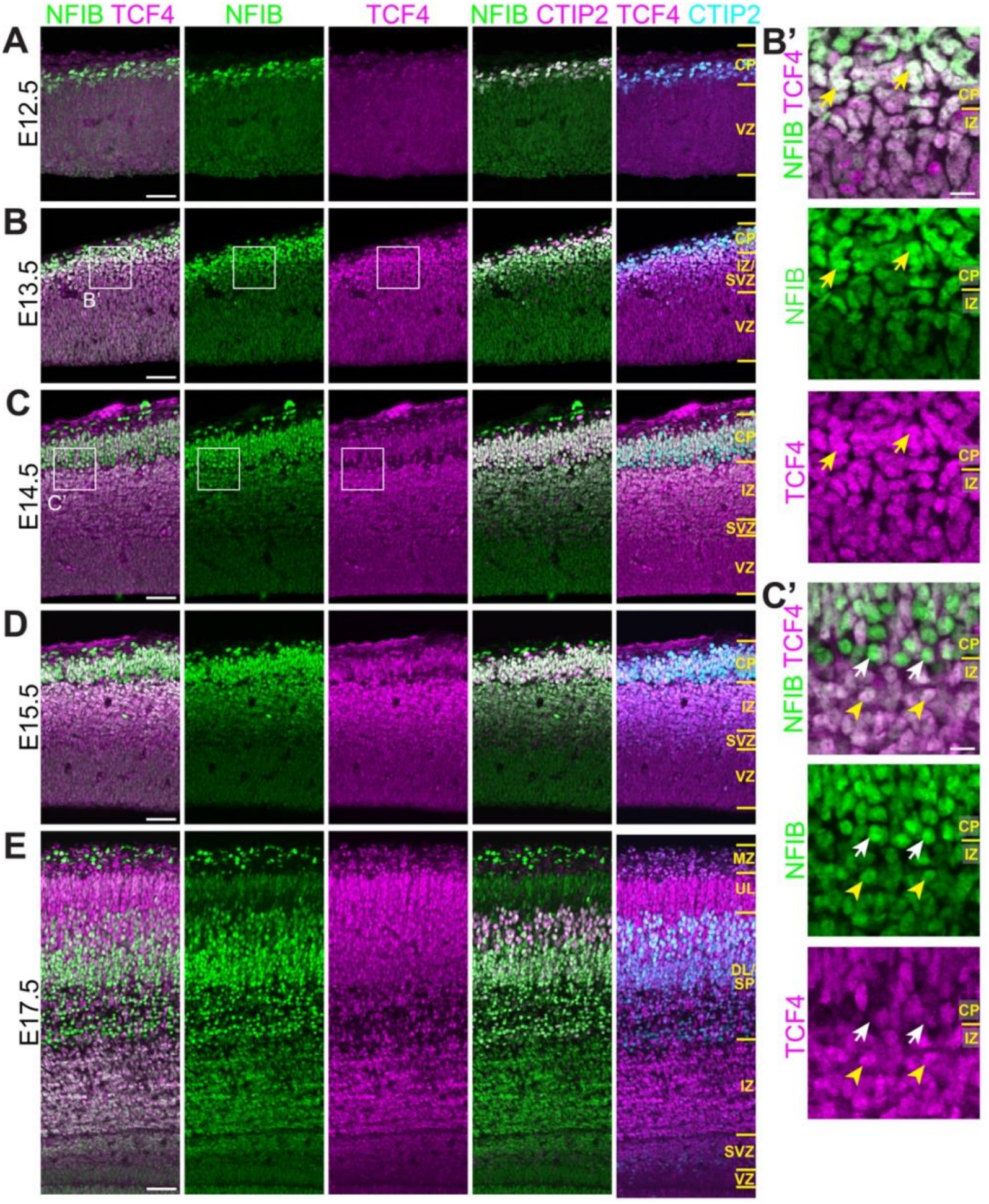
NFI and TCF4 are co-expressed in the cortical plate and ventricular zone during neocortical development. (*A*-*E*) Representative images of wildtype coronal sections between E12.5 and E17.5 stained for NFIB (green), TCF4 (magenta), and CTIP2 (pseudo-colored magenta or cyan to enable visualization). Insets in (*B*,*C*) represent high magnification images in (*B’*,*C’*). Yellow arrows in (*B’*) depict cells with high NFIB and TCF4 expression at E13.5. Yellow arrowheads in (*C’*) depict cells with high TCF4 but low NFIB expression. White arrows depict cells with high NFIB but low TCF4 expression. Scale bar, 50 µm for (*A*-*E*) and 10 µm for (*B’*,*C’*). (CP) Cortical plate, (DL) deeper layers, (IZ) intermediate zone, (MZ) marginal zone, (SP) subplate, (SVZ) subventricular zone, (UL) upper layers, (VZ) ventricular zone.

### NFIB and TCF4 are required for neuronal differentiation

Transcriptomic analyses of *Nfib* knockout neocortical tissue have been previously performed (Betancourt et al., 2014; Bunt et al., 2017). However, previous analyses were done at timepoints when changes in cell type composition because of delayed differentiation could result in indirect changes in gene expression (Betancourt et al., 2014; Piper et al., 2014). To identify direct targets of NFIB, we performed RNA-sequencing using E13.5 *Nfib* knockout embryos. We reasoned that changes in gene expression observed at this time point are more likely to represent *bona fide* targets. RNA-sequencing yielded 531 upregulated genes and 636 downregulated genes in *Nfib* knockout mice compared to wildtype littermates (Fig. 2A). Gene ontology of upregulated genes revealed enriched biological process terms broadly associated with development and neurogenesis (Table S1). This includes terms associated with progenitor self-renewal, such as ‘regulation of cell population proliferation’ (94 genes; FDR = 6.72e-7) and ‘regulation of Wnt signaling pathway’ (35 genes; FDR = 1.31e-6), which may underlie the expansion of the ventricular zone in these mice (Piper et al., 2014). Consistent with previous reports (Betancourt et al., 2014; Bunt et al., 2017), the top enriched terms associated with downregulated genes are linked with neuronal differentiation, such as ‘axon development’ (58 genes; FDR = 1.04e-17) and ‘synapse organization’ (53 genes; FDR = 7.6e-17) (Table S2). Therefore, changes in gene expression observed in the E13.5 *Nfib* knockout neocortex are consistent with delayed differentiation and cell type composition changes reported at later ages.

**Figure 2.**
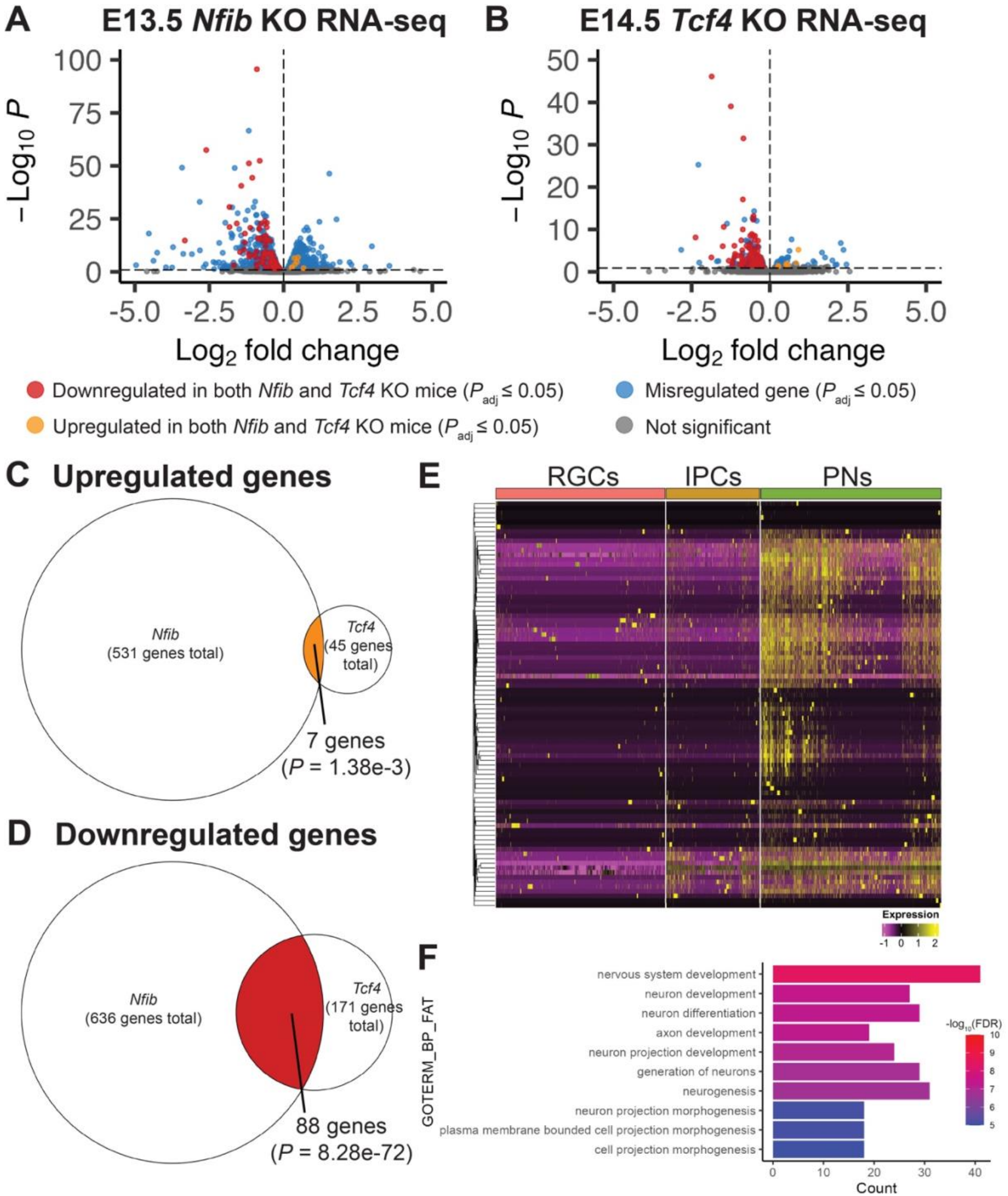
NFIB and TCF4 are required for the expression of a common set of 88 genes that are predominantly expressed in postmitotic neurons. (*A,B*) Volcano plots depicting differentially expressed genes in E13.5 *Nfib* knockout neocortex (*A*) and E14.5 *Tcf4* knockout neocortex (*B*). Shared upregulated genes are highlighted orange whereas shared downregulated genes are highlighted red. All other differentially expressed genes are depicted in blue. (*C*,*D*) Venn diagrams of overlapping up-(*C*) and downregulated genes (*D*) in *Nfib* and *Tcf4* knockout mice. *P* indicates hypergeometric distribution probability. (*E*) Clustered heatmap depicting relative expression of the 88 shared downregulated genes in the three major cell types within the E13.5 and E14.5 neocortex, obtained from a published single-cell dataset (Di Bella et al., 2021). Most genes are more highly expressed in postmitotic neurons as compared to the other cell types. Z-score normalization was independently applied to each gene across all cells and hierarchical clustering was performed based on Euclidean distance with complete linkage. The color scale represents z-score. Outliers were capped to the minimum and maximum values indicated. (RGCs) Radial glial cells, (IPCs) intermediate progenitor cells, (PNs) postmitotic excitatory neurons. (*F*) Enriched gene ontology terms associated with the 88 shared downregulated genes between *Nfib* and *Tcf4* knockout mice.

To compare our *Nfib* RNA-sequencing dataset with similar data from *Tcf4* knockout mice, we re-analyzed a published dataset generated using E14.5 *Tcf4* knockout neocortical tissue (Mesman et al., 2020). The original study reported six upregulated and 131 downregulated genes. When re-analyzing this data, we observed significant batch effects that would have reduced statistical power in the original analysis. We therefore accounted for this by normalization using RUVg (Risso et al., 2014) (Fig. S2). Using this approach but otherwise similar settings to the original analysis, we identified 45 upregulated genes and 171 downregulated genes (Fig. 2B). Four of the six upregulated genes in the original publication remained upregulated in our analysis. 99 of the 131 reported downregulated genes remained downregulated, suggesting high concordance between the two analysis strategies. Gene ontology on the 45 upregulated genes revealed only three enriched terms (Table S3). Consistent with the original analysis, downregulated genes were enriched for terms associated with neuronal differentiation (Table S4).

### NFIB and TCF4 regulate shared downstream targets

We compared both RNA-sequencing datasets for similarly misregulated genes. Seven genes were upregulated in both datasets (hypergeometric distribution *P* = 1.38e-3) (Fig. 2A-C and Table S5). Published data demonstrate that four of these, *Fgfr2*, *Prrx1*, *Sema5a*, and *Ticrr*, are preferentially expressed within the ventricular and subventricular zones at these stages (Fig. S3). Therefore, these genes could represent shared downstream targets of NFIB and TCF4 in radial glial cells and/or intermediate progenitors, where we observed lower expression of NFIB and TCF4 as compared to differentiating neurons.

Focusing next on shared downregulated genes, we identified 88 genes that were downregulated in both datasets (hypergeometric distribution *P* = 8.28e-72) (Fig. 2A,B,D and Table S6), many of which are predominantly expressed in postmitotic neurons (Fig. 2E and Table S7). Gene ontology demonstrates enrichment of biological process terms that were enriched amongst all downregulated genes from the individual datasets (Fig. 2F and Tables S2, S4, and S8). These terms include ‘nervous system development’, ‘neuron differentiation, and ‘axon development’, suggesting that while both NFIB and TCF4 regulate the same biological processes, part of their role occurs through the co-regulation of a common set of genes important for neuronal differentiation. Alternatively, given that we also identified many downregulated genes that were unique to each mouse model, NFIB and TCF4 may regulate distinct gene sets that are both critical for neuronal differentiation. In this instance, the 88 shared genes may represent convergence on a similar phenotype rather than shared direct targets. Indeed, 47 of the 88 genes demonstrated higher expression at E14.5 compared to E13.5 in ENCODE forebrain RNA-seq data (Table S9). Therefore, the 88 downregulated genes may be representative of delayed differentiation in both knockout mouse models rather than shared direct targets.

We also compared both datasets for discordantly misregulated genes (Table S10). Two genes were downregulated in the *Nfib* knockout neocortex but upregulated in the *Tcf4* knockout neocortex (hypergeometric distribution *P* = 0.276). An additional 12 genes were upregulated upon *Nfib* deletion but downregulated upon *Tcf4* deletion (hypergeometric distribution *P* = 0.019). Many of these genes are predominantly expressed in differentiating neurons (Fig. S4). Taken together with the 88 shared downregulated genes, this suggests that while NFIB and TCF4 may co-regulate a common set of genes important for neuronal differentiation, these transcription factors could also potentially function antagonistically in some contexts.

We further investigated the clinical relevance of shared genes by comparing these to high-confidence genes that were identified from a recent analysis of large-scale human cohorts (Fu et al., 2022). *Grin2b*, which was downregulated in both mouse models, is amongst 72 high-confidence genes identified from a cohort of 20,627 individuals with autism-spectrum disorder. *Fgfr2*, which was upregulated in both datasets, and eight of the 88 shared downregulated genes (*Cacna1e*, *Grin2b*, *Hivep2*, *Mef2c*, *Nfix*, *Sox5*, *Zbtb18*, and *Zeb2*) were amongst 309 high-confidence genes identified from a cohort of 31,058 individuals with severe developmental disorders (most of whom were diagnosed with intellectual disability).

As *NFIB* and *TCF4* haploinsufficiency are associated with CCD in both humans (Schanze et al., 2018; Zenker et al., 2019; Zollino et al., 2019; Zweier et al., 2007) and mice (Mesman et al., 2020; Piper et al., 2009a; Steele-Perkins et al., 2005), we also scrutinized shared misregulated genes for their association with CCD. Interestingly, five of the eight downregulated genes that are high-confidence genes for severe developmental disorders are associated with CCD in humans—*HIVEP2* (Srivastava et al., 2016; Zhu et al., 2026), *MEF2C* (Paciorkowski et al., 2013; Vrecar et al., 2017; Wan et al., 2021), *NFIX* (Priolo et al., 2018; Schanze et al., 2014; Shaw et al., 2010), *ZBTB18* (Cohen et al., 2017; Depienne et al., 2017; van der Schoot et al., 2018), and *ZEB2* (Amiel et al., 2001; Dastot-Le Moal et al., 2007; Zweier et al., 2002). Amongst the remaining shared downregulated genes, CCD is reported in humans with mutations affecting *AKT3* (Boland et al., 2007; Nakamura et al., 2014; Wang et al., 2025) and in *Akt3* knockout mice (Tschopp et al., 2005). Moreover, *Fezf2* (Chen et al., 2005), *Nrp1* (Lim et al., 2015; Piper et al., 2009b; Zhou et al., 2013), *Nuak1* (Courchet et al., 2013), *Sema7a* (Bessa et al., 2024), and *Shtn1* (Baba et al., 2018) are critical for callosal development, but no relevant phenotypes have been reported in humans. Taken together, these findings suggest that the overlapping set of genes that are misregulated in both mouse models may be important contributors to the shared phenotypes reported in *NFIB* and *TCF4* haploinsufficiency.

### Analogous genome-wide binding of NFIB and TCF4 in the developing neocortex

To further investigate the molecular convergence of NFIB and TCF4, we next utilized cleavage under targets and release using nuclease (CUT&RUN) to examine their genome-wide binding. NFIB and TCF4 CUT&RUN, performed using cells dissociated from E14.5 wildtype neocortical tissue, identified highly reproducible binding at 11,854 and 13,760 peaks, respectively. *De novo* motif discovery indicate that these peaks represent true binding sites, as the top enriched motif at NFIB peaks corresponded to the NFI half-site (Fig. 3A) or palindromic dimer motif (Fig. 3B) (Gronostajski, 1986; Gronostajski et al., 1985). The top enriched motif at TCF4-bound peaks was its cognate E-box motif (Fig. 3A,B). We also found enrichment of the NFI half-site and dimeric motifs at TCF4-bound peaks (NFI motifs at TCF4 peaks underlined in magenta in Fig. 3A).

**Figure 3.**
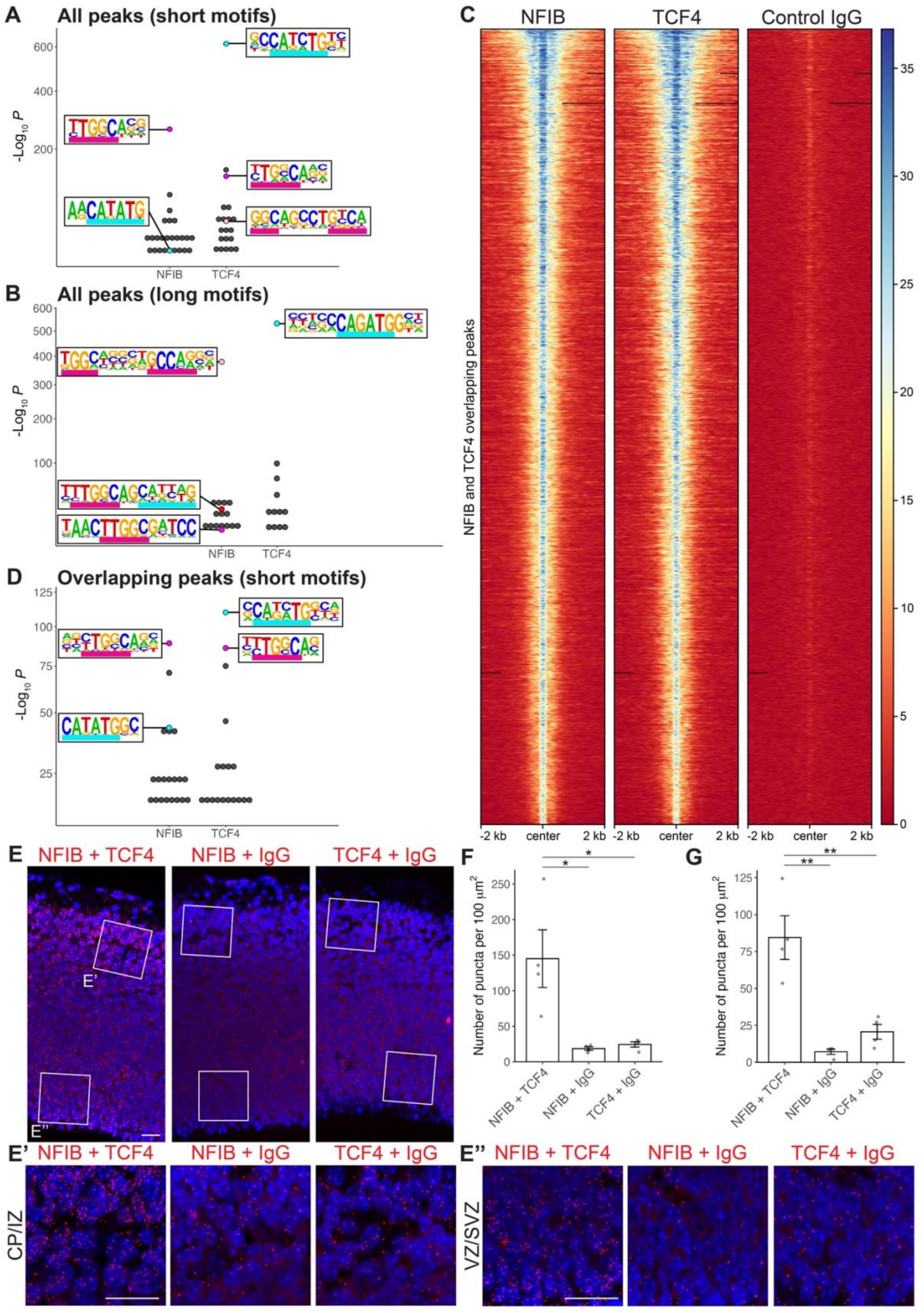
Genome-wide binding of NFIB and TCF4 in the E14.5 neocortex largely overlap each other. (*A*,*B*) Enriched motifs identified from NFIB and TCF4 CUT&RUN peaks. *De novo* motif discovery was performed to identify enriched motifs of 8 to 12 (*A*) or 14 to 16 (*B*) nucleotides in length. Horizontal magenta and cyan bars represent motifs resembling the NFI half-site and E-box motifs, respectively (also for *D*). The NFI dimeric motif is represented by two magenta bars separated by five degenerate nucleotides. (*C*) Heatmaps depicting fragment density at overlapping NFIB and TCF4 CUT&RUN peaks. Genomic regions were sorted by mean pileup of fragments across all samples. Black bars indicate regions void of any fragments. (*D*) Enriched motifs from overlapping NFIB and TCF4 CUT&RUN peaks. (*E*) Duolink PLA reveal potential interaction between NFIB and TCF4. Representative PLA images of E14.5 wildtype coronal sections, using antibodies specific for NFIB and TCF4 (red), or their corresponding isotype controls, and counterstained with DAPI (blue). Insets in (*E*) represent high magnification images in (*E’*,*E **’***). Scale bars, 20 µm. (CP) Cortical plate, (IZ) intermediate zone, (SVZ) subventricular zone, (VZ) ventricular zone. (*F*,*G*) Quantification of PLA puncta within the CP/IZ (*F*) and VZ/SVZ (*G*). Data represents mean ± s.e. with individual points representing biological replicates. Two-tailed Student’s t-tests: (*) *P* = 0.021 for NFIB + TCF4 versus NFIB + IgG, 0.025 for NFIB + TCF4 versus TCF4 + IgG in (*F*), (**) *P* = 0.002 for NFIB + TCF4 versus NFIB + IgG, 0.006 for NFIB + TCF4 versus TCF4 + IgG in (*G*).

Additionally, we discovered an enriched motif within NFIB-bound peaks that resembled a combination of an NFI half site and E-box motif occurring adjacent to each other (NFI half-site motif underlined in magenta and E-box motif underlined in cyan in Fig. 3B). This suggests that NFIB and TCF4 could regulate an overlapping set of genes by binding to their cognate motifs within the same regulatory elements. Collectively, these analyses also demonstrated that our CUT&RUN experiments were successful in identifying genome-wide binding of these transcription factors.

Given our observation of a motif resembling a combination of the NFI half site and E-box motifs, we sought to investigate the co-enrichment of these motifs as compared to other transcription factor motifs. To do this, we examined motif co-occurrences at all NFIB and TCF4 peaks using TF-COMB (Bentsen et al., 2022). The NFI and E-box motifs pair was amongst the top co-enriched transcription factor pair within both datasets (Fig. S5). Other top pairs include a motif recognized by the Notch effector RBPJ that was co-enriched with both the NFIB and TCF4 motifs and a motif recognized by MEIS1/2 with the TCF4 motif. RBPJ and MEIS2 were recently demonstrated to function cooperatively with NFIB and TCF4 in other cellular contexts (Bright et al., 2025; Le et al., 2024). Therefore, our findings suggest that these previous findings may also extend to neocortical development. More importantly, these analyses demonstrate that the NFIB and TCF4 motifs are co-enriched within regulatory elements bound by these transcription factors in the developing neocortex. This suggests that NFIB and TCF4 could regulate the same target genes by each binding to their own cognate motifs within the same regulatory elements.

We next compared both CUT&RUN datasets and identified 6,230 NFIB-bound peaks (52.6% of all NFIB peaks) that overlapped 6,236 TCF4-bound peaks (45.3% of all TCF4 peaks) (Fig. 3C). We repeated *de novo* motif discovery on overlapping peaks and found strong enrichment of both the NFI half-site and E-box motifs (Fig. 3D). Co-binding of NFIB and TCF4 to the same regulatory elements is also supported by proximity ligation assays (PLAs). We performed PLAs on E14.5 wildtype sections using NFIB and TCF4 antibodies in combination with one another or with negative control mouse or rabbit IgG, respectively (Fig. 3E). We observed a significant increase in the number of PLA puncta with NFIB and TCF4 as compared to negative control conditions, suggesting their close co-localization within individual cells (Fig. 3F,G). We also observed more PLA puncta within the cortical plate and intermediate zone as compared to the ventricular/subventricular zone, consistent with higher NFIB and TCF4 expression in this region. Taken together with our CUT&RUN data, this suggests that NFIB and TCF4 regulate an overlapping set of genes, at least in part, by co-binding to the same regulatory elements.

We also investigated non-overlapping CUT&RUN peaks by examining the density of TCF4-bound fragments around NFIB-specific peaks and vice versa. These analyses demonstrate that both NFIB and TCF4 may still bind to many non-overlapping peaks, albeit with weaker binding observed for one transcription factor (Fig. S6). This weaker binding did not pass the stringent cutoff used for peak calling, resulting in such peaks being called for only one transcription factor and not the other.

### NFIB and TCF4 binding are associated with active promoters and distal enhancers

We next annotated overlapping and non-overlapping peaks, taking into consideration whether each peak was located proximally (defined as within two kilobases of a transcription start site) or distally to its nearest gene. We annotated peaks by comparing each peak set to ENCODE E14.5 ChromHMM chromatin states derived from forebrain tissue (Gorkin et al., 2020). Overlapping and unique proximal peaks were strongly associated with active followed by weak promoter states (Pr-A and Pr-W, respectively, in Fig. S7A), which are defined by high H3K4me3 and differential H3K27ac enrichment (Fig. S8A). Overlapping distal peaks were associated with strong and poised distal enhancer states (En-Sd and En-Pd in Fig. S7B), which are enriched for H3K27ac and H3K4me1 (Fig. S8B). Non-overlapping NFIB and TCF4 distal peaks showed weaker association with the strong distal enhancer state but pronounced association with other chromatin states, such as the transcription permissive state or chromatin regions with inadequate signal (Tr-P and NS in Fig. S7B). Therefore, peak annotation suggests that overlapping NFIB and TCF4 peaks are more likely to correspond to active regulatory elements as compared to peaks bound by NFIB or TCF4 alone. Overlapping NFIB and TCF4 peaks were also negatively correlated with repressive chromatin states and their underlying histone modifications (Hc-P and Hc-H in Fig. 7 and Fig. S8C,D). This suggests that NFIB and TCF4 bind to a common set of active promoters and distal enhancers during neocortical development, presumably to drive expression of the genes associated with these regulatory elements.

### Binding of NFIB and TCF4 is associated with transcriptional activation

To examine how NFIB and TCF4 binding affects gene regulation, we integrated our CUT&RUN data with both RNA-sequencing datasets. To do this, we first assigned proximal binding of NFIB and TCF4 to their nearest gene. Next, we used ENCODE predicted enhancer targets inferred using two independent methods to assign distal peaks to putative target genes (Gorkin et al., 2020; Yu et al., 2025). 41.6% and 38.5% of NFIB and TCF4 distal peaks, respectively, were assigned to at least one target gene. Finally, all remaining peaks were assigned to their nearest gene, resulting in almost all NFIB and TCF4 binding sites assigned to at least one gene (Tables S11 and S12). NFIB and TCF4 target genes were then intersected with the RNA-sequencing datasets to identify differentially expressed genes that were directly bound by NFIB or TCF4.

NFIB and TCF4 binding were detected at 45.1% and 43.8% of all gene loci, respectively, with many of these being direct binding to the promoter (Fig. S9). In line with studies on other transcription factors (Kribelbauer et al., 2019; MacQuarrie et al., 2011) most binding events are not associated with differentially expressed genes upon *Nfib* or *Tcf4* deletion.

Focusing on differentially expressed genes from the *Nfib* knockout neocortex, we observed that a larger proportion of these genes were direct targets of NFIB as compared to all genes (Fig. S9A). This increase could be attributed to NFIB binding particularly at distal sites, which presumably function as enhancers to regulate gene expression. We quantified the enrichment of NFIB binding to both up- and downregulated genes using permutation tests, which repeatedly samples an equal number of expression-matched control genes across 10,000 permutations. We found that NFIB binding to both up- and downregulated genes is significantly enriched compared to permuted controls (Fig. 4A; *P* < 1e-5 in all instances). Importantly, we observed the highest fold enrichment with the 88 shared downregulated genes. 73 of the 88 genes were direct targets of NFIB. 60 of these genes demonstrated binding at the promoter and/or distal binding inferred from the integrated datasets (Gorkin et al., 2020; Yu et al., 2025) (Fig. S9A). These findings suggest that NFIB primarily functions as a transcriptional activator during neocortical development, but that NFIB could also repress a subset of its downstream targets upon binding to their regulatory elements.

**Figure 4.**
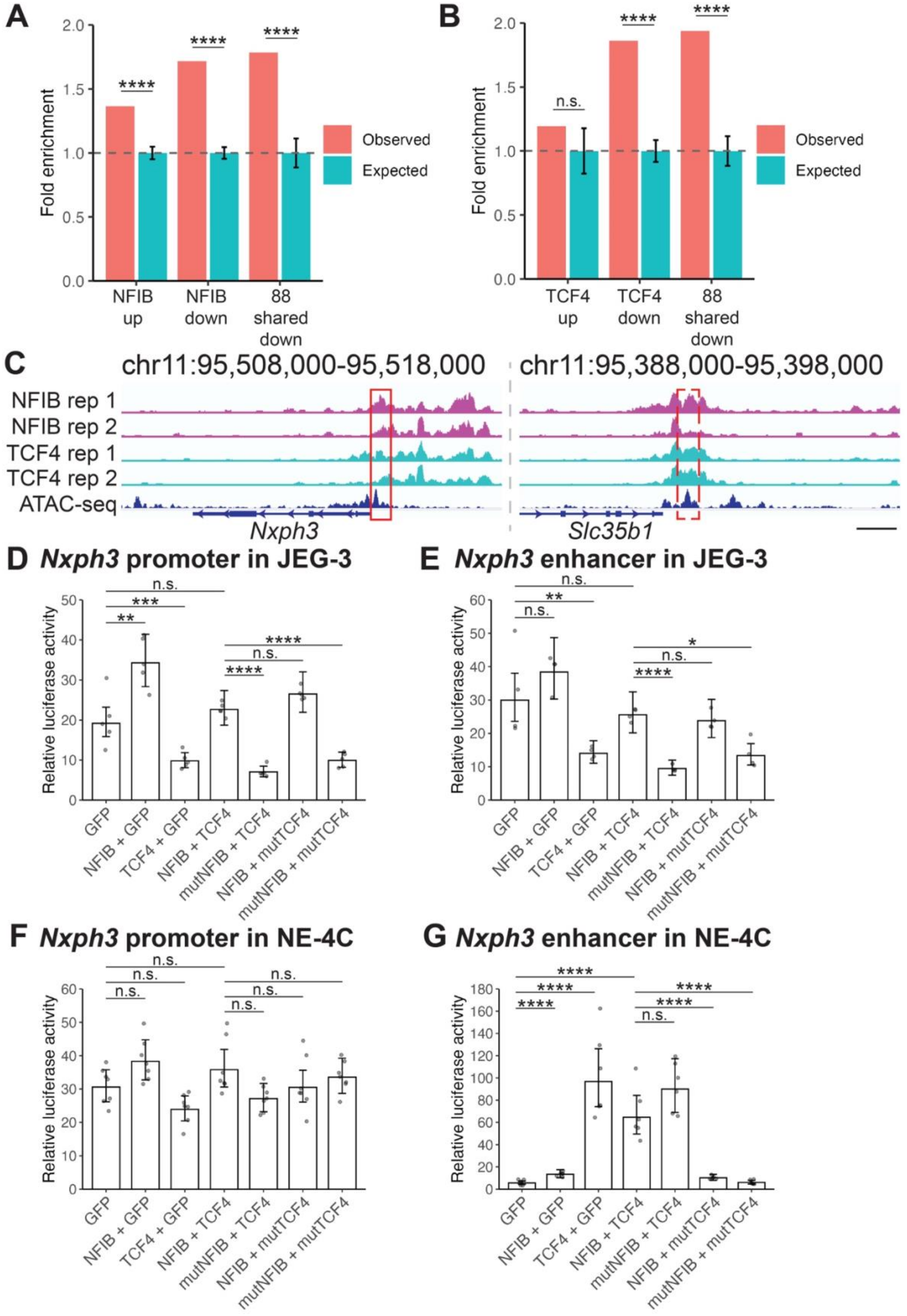
NFIB and TCF4 co-expression in luciferase assays elicit combinatorial effects on transcriptional activity. (*A*,*B*) Downregulated genes in the *Nfib* and *Tcf4* knockout neocortices are significantly enriched for NFIB and TCF4 binding, respectively. Bar charts depict fold enrichment compared to expected enrichment derived from 10,000 permutations of randomly sampled expression-matched control genes. Expected bars represent mean ± s.d. from 10,000 permutations. Permutation tests in (*A*,*B*): (****) *P* < 1e-5, (n.s.) *P* = 0.172. (C) Genome browser plots demonstrating NFIB and TCF4 binding at the *Nxph3* promoter (left panel) and an ENCODE-predicted distal enhancer (right panel). The ATAC-seq track represents ENCODE E14.5 forebrain ATAC-seq data (Gorkin et al., 2020). The red boxes represent the promoter (solid line) and enhancer (dashed line) regions cloned for luciferase assays. (*D*-*G*) Relative luciferase activities of the *Nxph3* promoter (*D*,*F*) and distal enhancer (*E*,*G*) upon co-expression with wildtype or missense variants of NFIB and TCF4 in JEG-3 (*D*,*E*) and NE-4C (*F*,*G*) cells. Data represents back-transformed geometric mean ± 95% c.i. Individual points represent biological replicates adjusted for batch by adding each observation’s residual from the ANOVA model to the estimated marginal mean and back-transforming. Two-tailed Student’s t-test in (*D*): (**) *P* = 0.0005, (***) *P* = 0.0001, (****) *P* < 0.0001, (n.s.) *P* = 0.4302. Two-tailed Student’s t-test in (*E*): (*) *P* = 0.0029, (**) *P* = 0.0008, (****) *P* < 0.0001, (n.s.) *P* = 0.4185 for GFP versus NFIB + GFP, 0.6706 for GFP versus NFIB + TCF4 and NFIB + TCF4 versus NFIB + mutTCF4. Two-tailed Student’s t-test in (*F*): (n.s.) *P* = 0.1886 for GFP versus NFIB + GFP, 0.1461 for GFP versus TCF4 + GFP, 0.4494 for GFP versus NFIB + TCF4 and NFIB + TCF4 versus NFIB + mutTCF4, 0.0907 for NFIB + TCF4 versus mutNFIB + TCF4, and 0.5569 for NFIB + TCF4 versus mutNFIB + mutTCF4. Two-tailed Student’s t-test in (*G*): (****) *P* < 0.0001, (n.s.) *P* = 0.0823.

Performing identical analyses on TCF4 target genes, we found significant enrichment of TCF4 binding to downregulated but not upregulated genes (Fig. 4B; *P* = 0.172 for upregulated genes and < 1e-5 for downregulated genes). Again, the 88 shared downregulated genes demonstrated the highest fold enrichment, with 77 of these being direct TCF4 targets (*P* < 1e-5). 72 of these demonstrated binding at the promoter and/or distal binding inferred from the integrated datasets (Gorkin et al., 2020; Yu et al., 2025) (Fig. S9B). Therefore, like NFIB, TCF4 likely functions primarily as a transcriptional activator. Furthermore, direct binding of NFIB and TCF4 to the majority of the 88 shared downregulated genes, including all eight high-confidence genes implicated in severe human developmental disorders and the 11 genes associated with corpus callosum development (Tables S11 and S12), demonstrate that NFIB and TCF4 molecularly converge by binding to the same regulatory elements to regulate the expression of a common set of genes.

### Differential transcriptional activation upon NFIB and TCF4 binding to the same regulatory elements

To further study the molecular convergence of NFIB and TCF4 during neocortical development, we next sought to determine the effect of NFIB and TCF4 co-expression on individual regulatory elements. We examined this using *in vitro* luciferase reporter assays, selecting the promoter and a distal enhancer of *Nxph3* as candidate regulatory elements for analysis. *Nxph3* is a neuronal-specific gene that is strongly reduced upon *Nfib* and *Tcf4* deletion (log_2_ fold change = -1.82, *P*_adj_ = 7.69e-22 for *Nfib* knockout; log_2_ fold change = -1.5, *P*_adj_ = 9.06e-7 for *Tcf4* knockout). Additionally, CUT&RUN demonstrates that NFIB and TCF4 bind both these regulatory elements (Fig. 4C). *Nxph3* is specifically expressed in human (Velmeshev et al., 2023) and mouse layer 6 neurons and preplate cells during prenatal development (Fig. S10). *Nxph3* knockout mice display no obvious structural brain deficits but exhibit abnormalities in sensorimotor gating and motor coordination (Beglopoulos et al., 2005), demonstrating its importance for neuronal function. While NXPH3 has not been implicated in a NDD, it functions as a ligand that binds strongly to NRXN1 at synapses (Missler et al., 1998; Petrenko et al., 1993; Petrenko et al., 1996). Loss-of-function of both copies of *NRXN1* in human results in a NDD known as Pitt-Hopkins-like syndrome 2. As its namesake suggests, this NDD bears many similarities to *TCF4* haploinsufficiency (Harrison et al., 2011; Zweier et al., 2009). Unlike *Nxph3* however, *Nrxn1* is discordantly regulated (log_2_ fold change = -0.3, *P*_adj_ = 0.023 in *Tcf4* knockout mice and log_2_ fold change = 0.19, *P*_adj_ = 0.048 in *Nfib* knockout mice).

We independently cloned the *Nxph3* promoter and distal enhancer to drive expression of firefly luciferase and transfected these into human JEG-3 cells or mouse neuroepithelial NE-4C cells. JEG-3 cells were selected as they lowly express NFI transcription factors (Bruggemeier et al., 1990; Chaudhry et al., 1998) (Fig. S11), whereas NE-4C cells provided a context comparable to cortical development. We tested each reporter with co-expression of control GFP, wildtype NFIB, wildtype TCF4, or NFIB and TCF4 combined. NFIB activated the *Nxph3* promoter in JEG-3 cells and the distal enhancer in NE-4C cells (1.9-fold increase for NFIB + GFP versus control GFP in Fig. 4D and 2.4-fold increase in Fig. 4G; *P* = 0.005 and < 0.0001, respectively). We observed a similar trend of NFIB-mediated activation of the enhancer in JEG-3 cells (Fig. 4E) and the promoter in NE-4C cells (Fig. 4F), but neither were statistically significant (*P* = 0.4185 and 0.1886, respectively). Contrary to NFIB, TCF4 repressed the promoter and distal enhancer in JEG-3 cells (1.9-fold decrease for TCF4 + GFP versus control GFP in Fig. 4D and 2.1-fold decrease in Fig. 4E; *P* = 0.0001 and 0.0008, respectively). A similar albeit statistically insignificant trend towards repression was observed for the promoter in NE-4C cells (Fig. 4F; *P* = 0.1461). In contrast to this and what we observed in JEG-3 cells, TCF4 strongly activated the *Nxph3* enhancer in NE-4C cells (17.3-fold increase for TCF4 + GFP versus GFP alone in Fig. 4G; *P* < 0.0001). This demonstrates that TCF4-mediated activation of the *Nxph3* enhancer depends on cellular context.

When we co-expressed NFIB and TCF4 in JEG-3 cells, we observed an intermediate level of activity between that of NFIB and TCF4 by themselves. In both scenarios where we previously observed activation by NFIB and repression by TCF4, co-expression of both resulted in an intermediate level of activity that was not significantly different from GFP control (*P* = 0.4302 for NFIB + TCF4 versus control GFP in Fig. 4D and 0.6706 in Fig. 4E). This reinforces the opposite roles that NFIB and TCF4 play in regulating transcriptional activity in JEG-3 cells. We observed a similar result for the *Nxph3* enhancer that was activated by NFIB and TCF4 in NE-4C cells. In this instance, we observed an 11.5-fold increase in activity compared to GFP control, between that of NFIB and TCF4 by themselves (Fig. 4G; *P* < 0.0001). Therefore, these findings suggest that NFIB and TCF4 function in a competitive manner to regulate the *Nxph3* promoter and enhancer.

We also performed luciferase assays to investigate whether NFIB and TCF4 regulate *Sstr2* promoter activity. *Sstr2* expression was significantly reduced in our E13.5 *Nfib* knockout RNA-seq data (log_2_ fold change = -0.44, *P*_adj_ = 7.8e-11) but not in the E14.5 *Tcf4* knockout dataset (log_2_ fold change = -0.32, *P*_adj_ = 0.247). *Sstr2* encodes somatostatin receptor 2, which is expressed in a subset of intermediate progenitors and postmitotic neurons during neocortical development and regulates cell migration and axonal outgrowth (Bedogni and Hevner, 2021; Le Verche et al., 2009). *Sstr2* is a known target of TCF4. TCF4 activates *Sstr2* expression by binding to a regulatory element within an intron that was previously thought to be the promoter region (Dorflinger et al., 1999). Since we did not detect NFIB or TCF4 binding at this intronic region (Fig. S12A), we focused instead on the actual promoter, which has not been previously examined. Luciferase assays in JEG-3 cells yielded similar results to the *Nxph3* promoter and distal enhancer in this cell line (Fig. S12B). We observed a 2.4-fold increase in activity with NFIB + GFP compared to control GFP (*P* = 0.043). TCF4 repressed promoter activity by 1.9-fold (*P* = 0.025). Again, luciferase activity upon co-expression of NFIB and TCF4 was not significantly different from control GFP (*P* = 0.7546), reinforcing the notion that NFIB and TCF4 can function antagonistically to regulate transcriptional activity.

We next examined how co-expression of missense variants of NFIB (p.Lys126Glu) and TCF4 (p.Arg580Trp) affected luciferase activity, comparing these to co-expression of wildtype NFIB and TCF4. These variants were discovered from individuals with *NFIB* (Schanze et al., 2018) or *TCF4* (Amiel et al., 2007; Zweier et al., 2007) haploinsufficiency. Both missense variants occur within the DNA-binding domains of the respective proteins and was demonstrated to disrupt DNA binding for TCF4 (Sepp et al., 2012). In previous luciferase assays, both missense variants demonstrate reduced activity compared to their wildtype counterparts (Forrest et al., 2012; Schanze et al., 2018; Sepp et al., 2012; Zweier et al., 2007). Replacing wildtype NFIB with mutant NFIB in JEG-3 cells abolishes transcriptional activation, resulting in repression when co-expressed with wildtype TCF4 (mutNFIB + TCF4 versus NFIB + TCF4 in Fig. 4D,E, and Fig. S12B). We observed a reduction of 2.8-fold, 2.7-fold, and 2.5-fold at the *Nxph3* promoter, *Nxph3* enhancer, and *Sstr2* promoter, respectively (*P* < 0.0001 for the *Nxph3* promoter and enhancer, 0.0031 for the *Sstr2* promoter). A similar albeit statistically insignificant effect was observed for the *Nxph3* enhancer in NE-4C cells. In this instance, replacing wildtype NFIB with mutant NFIB results in increased activity when co-expressed with wildtype TCF4, which strongly activates this enhancer (1.4-fold increase; *P* = 0.0823). Therefore, our results demonstrate that DNA-binding is critical for NFIB-mediated activation of these regulatory elements.

On the contrary, replacing wildtype TCF4 with its missense variant resulted in a significant change in activity only at the *Nxph3* enhancer in NE-4C cells (Fig. 4G). Compared with co-expression of wildtype NFIB and TCF4, mutant TCF4 reduced transcriptional activity by 6.3- fold when co-expressed with wildtype NFIB and 10.6-fold when co-expressed with mutant NFIB (*P* < 0.0001 in both instances). Taken together with our findings from co-expressing mutant NFIB with wildtype TCF4, this indicates that NFIB and TCF4 regulate *Nxph3* enhancer activity in NE-4C cells by binding competitively to this region.

In JEG-3 cells, the TCF4 missense variant retained its ability to repress transcriptional activity. We found no significant differences in relative luciferase activity between co-expression of mutant TCF4 plus wildtype NFIB and wildtype TCF4 plus NFIB across all three regions (*P* = 0. 4302, 0.6706, and 0.69 for NFIB + mutTCF4 versus NFIB + TCF4 at the *Nxph3* promoter, enhancer, and *Sstr2* promoter, respectively). Repression by mutant TCF4 was similarly observed when we co-expressed both missense variants (mutNFIB + mutTCF4). In this instance, we observed a reduction in activity consistent with reduced activation by mutant NFIB (1.8-fold, 1.9-fold, and 2-fold reduction for the *Nxph3* promoter, enhancer, and *Sstr2* promoter, respectively; *P* < 0.0001, *P* = 0.0029 , and *P* = 0.0151, respectively). These findings indicate that NFIB and TCF4 do not function competitively in our luciferase assays in JEG-3 cells. Rather, repression by TCF4 is independent of its ability to bind to DNA and could be mediated by one of the following scenarios. First, TCF4 may be indirectly recruited through its interaction with an unidentified transcriptional regulator that binds to these sites. Second, over-expression of TCF4 may result in an excess of TCF4 protein that does not bind to DNA but retains its ability to interact with other transcriptional regulators/machinery. In this case, TCF4 could sequester these proteins thereby reducing transcriptional activity. In both scenarios, TCF4 represses activity without directly binding to DNA.

In summary, our luciferase assays demonstrate that NFIB and TCF4 expression can lead to different transcriptional outcomes at individual regulatory elements, with their co-expression resulting in a combinatorial effect on transcriptional activity. NFIB binds to both the *Nxph3* promoter and enhancer to drive their activation in JEG-3 cells. NFIB similarly activates the *Nxph3* enhancer in NE-4C cells. On the contrary, TCF4 indirectly represses *Nxph3* promoter and enhancer activity in JEG-3 cells but functions as a potent activator of the *Nxph3* enhancer in NE-4C cells. Furthermore, NFIB and TCF4 compete to bind to and regulate this enhancer in NE-4C cells. We propose that the antagonistic functions of NFIB and TCF4 across different regulatory elements may be a means through which gene expression is fine-tuned to ensure appropriate levels during neocortical development.

## Discussion

In this study, we investigated whether NFIB and TCF4 molecularly converge by binding to the same cis-regulatory elements in mice. Previous studies that performed chromatin immunoprecipitation and high-throughput sequencing (ChIP-seq) using antibodies that recognize multiple (or other) genes belonging to these transcription factor families have demonstrated the co-enrichment of their binding sites within distal enhancers of neural cells (Mateo et al., 2015; Quevedo et al., 2019). Our analyses expand upon previous work by revealing remarkable overlap in the genome-wide binding of NFIB and TCF4 during neocortical development. Consistent with their shared binding at active regulatory elements, we also demonstrate that NFIB and TCF4 co-regulate an overlapping set of genes in differentiating neurons. Despite previous data demonstrating that NFIB is highly expressed in differentiating neurons (Betancourt et al., 2014; Bunt et al., 2015; Bunt et al., 2017; Plachez et al., 2008), its function in this cell type is often overlooked due to its prominent role as a gliogenic factor (Caiazzo et al., 2015; Deneen et al., 2006; Yeon et al., 2021). Critically, our findings demonstrate that neurogenesis is not merely delayed in the absence of NFIB (Betancourt et al., 2014; Bunt et al., 2017), but that NFIB directly regulates genes that are important for neuronal differentiation.

Taken together with previous studies, our CUT&RUN data and PLAs suggest that these transcription factors interact with one another and with other transcriptional co-regulators (Bright et al., 2025; Malaymar Pinar et al., 2025; Martins-Costa et al., 2024; Moen et al., 2017; Quevedo et al., 2019). Part of their role in regulating gene expression is recruitment of the Mediator complex (Quevedo et al., 2019). NFIB and TCF4 also interact with the SWI/SNF complex to regulate chromatin accessibility (Malaymar Pinar et al., 2025; Martins-Costa et al., 2024). We speculate that this function enables the binding of other key transcription factors required for target gene expression. This may explain why *Nxph3*, which requires both NFIB and TCF4 for its expression, is not expressed in all cells that co-express NFIB and TCF4 but is confined to specific cortical layers (Fig. S10) (Beglopoulos et al., 2005; Velmeshev et al., 2023). Future studies investigating how loss of either NFIB or TCF4 affects the binding of each other as well as the binding of other interacting proteins could further uncover how these proteins function.

We identified 88 shared downregulated genes between the two RNA-seq datasets. The majority of these were direct targets of NFIB and TCF4 (Tables S11 and S12). Recent studies demonstrate that NFIB and TCF4 may also co-regulate overlapping sets of target genes during the development of mouse cortical interneurons and human callosal neurons (Bright et al., 2025; Martins-Costa et al., 2024). We propose that shared target genes, especially the eight that are recognized as high-confidence genes implicated in severe developmental disorders in humans and the 11 genes associated with CCD in humans and/or mice, could serve as high priority targets to further understand the neurological phenotypes that are common to both *NFIB* and *TCF4* haploinsufficiency. Furthermore, of the eight genes implicated in severe developmental disorders, six are transcription factors (*Hivep2*, *Mef2c*, *Nfix*, *Sox5*, *Zbtb18*, and *Zeb2*) whereas the remaining two are involved in synaptic function (*Cacna1e* and *Grin2b*). Hence, all eight genes converge onto two functional pathways (transcriptional regulation and synaptic function) commonly associated with NDDs (Hormozdiari et al., 2015; Parenti et al., 2020) and may regulate overlapping cellular and molecular mechanisms.

Previous functional studies on four (*Nfix*, *Hivep2*, *Mef2c*, and *Zbtb18*) of the six high-confidence NDD target genes that encode for transcription factors have uncovered additional links to NFIB or TCF4. We previously demonstrated that NFIB transcriptionally activates *Nfix* in the developing spinal cord (Matuzelski et al., 2017). Moreover, NFIX binds to the same binding sites as NFIB (Fraser et al., 2020), potentially as heterodimers (Kruse and Sippel, 1994). In a similar manner, HIVEP2 and TCF4 is thought to function cooperatively in transactivating *Sstr2* expression in cortical glutamatergic neurons. A previous study inferred from examining expression patterns that *Sstr2* is only expressed in cells that co-express TCF4 and HIVEP2 (Dorflinger et al., 1999). This is supported by luciferase assays that demonstrated that TCF4 and HIVEP2 binding sites within an *Sstr2* intronic regulatory element were required for its activity.

However, *Sstr2* was not differentially expressed in our analysis of the *Tcf4* knockout RNA-seq data. Furthermore, we did not observe TCF4 binding to this intronic region but instead detected NFIB and TCF4 binding to the actual promoter. Interestingly, our luciferase assays in JEG-3 cells demonstrate that NFIB activates the promoter but TCF4 represses it. Further tests are thus required to determine whether TCF4 activates *Sstr2* expression through another regulatory element. Aside from this, *MEF2C* haploinsufficiency results in diminished *MECP2* and *CDKL5* expression (Zweier et al., 2010). MEF2C was found to function synergistically with an unidentified transcription factor that binds to E-box motifs to regulate these genes. Our CUT&RUN data demonstrates that TCF4 binds to the *Mecp2* and *Cdkl5* promoters, but neither were differentially expressed in *Tcf4* knockout mice. The transcriptional repressor ZBTB18 also binds to E-box motifs (Aoki et al., 1998). ZBTB18 binds to this motif within the *Neurod1* promoter to repress its expression (Xiang et al., 2012). NEUROD1 is critical for neurogenesis and can heterodimerize with TCF4 (Forrest et al., 2012). Interestingly, we did not detect TCF4 binding to the *Neurod1* promoter but *Neurod1* expression is upregulated in the *Tcf4* knockout RNA-seq dataset. Therefore, ZBTB18 may function in a negative feedback loop with TCF4 by competing for the same binding sites such as in the *Neurod1* promoter while simultaneously repressing the expression of TCF4’s heterodimerization partners. Taken together, these studies demonstrate that NFIB and TCF4 form complex regulatory relationships with the downstream transcription factors that they regulate. Future work aiming to further understand the underlying etiology of shared phenotypes could investigate molecular convergence between these transcription factors and NFIB or TCF4.

We observed that NFIB and TCF4 also bind to regulatory elements within the *Nfib* and *Tcf4* loci (Fig. S13). Therefore, both transcription factors function in a co-regulatory feedback loop as proposed previously (Quevedo et al., 2019). In support of this, *Nfib* is downregulated in the *Tcf4* knockout neocortex (log_2_ fold change = -0.36; *P*_adj_ = 1.40e-6). This was also observed in E18.5 *Tcf4* knockout *Satb2*-positive cortical neurons (Wittmann et al., 2021) and in neural stem cells upon *Tcf4* knockdown (Quevedo et al., 2019). Recent luciferase assays demonstrate that NFIB also drive its own expression (Zhang et al., 2026). Therefore, TCF4 and NFIB both regulate *Nfib* expression. In contrast to this, *Tcf4* expression is unchanged upon *Nfib* deletion despite NFIB binding to the *Tcf4* locus. While NFIB might regulate *Tcf4* in a different context (Bright et al., 2025), we did not observe *Tcf4* downregulation in our E13.5 or E16.5 *Nfib* knockout RNA-seq datasets (this study and Bunt et al., 2017, respectively). This raises an important point for consideration: NFIB and TCF4 bind to many genes that were not differentially expressed upon deletion of these proteins. This may be explained in part by compensation by other *Nfi* or *Tcf* genes. Like NFIB, NFIA and NFIX are expressed in radial glia and postmitotic neurons during neocortical development (Campbell et al., 2008; Plachez et al., 2008). Previous studies demonstrate that concurrent deletion of two *Nfi* genes results in more severe brain phenotypes as compared to deletion of a single *Nfi* gene, indicating that they have overlapping but non-redundant functions in neocortical development (Bunt et al., 2017; Zhang et al., 2026). NFIA and NFIX also bind to many of the same regulatory elements as NFIB (Fraser et al., 2020), potentially as NFI heterodimers (Bunt et al., 2017; Chaudhry et al., 1998; Kruse and Sippel, 1994). Furthermore, haploinsufficiency of *NFIA* or *NFIX* in humans results in the same neurological phenotypes observed with *NFIB* haploinsufficiency (Zenker et al., 2019). Therefore, NFIA and NFIX may compensate for NFIB in regulating *Tcf4* expression as well as other genes. Similarly, TCF3 and TCF12 can compensate for one another as well as TCF4, as phenotypes that are absent in single knockouts become apparent upon deletion of two of these genes (Ravanpay and Olson, 2008; Talley et al., 2023). In humans, neurological phenotypes such as developmental delay and CCD that are observed in Pitt-Hopkins syndrome are also reported in individuals with *TCF12* mutations (Paumard-Hernandez et al., 2015; Sharma et al., 2013). Therefore, functional redundancy between different members of each transcription factor family may moderate phenotypes that are observed with deletion of multiple alleles of each family member.

While both NFIB and TCF4 transactivate *Nxph3*, our luciferase assays indicate that they bring about vastly different outcomes at the promoter and enhancer elements. NFIB directly activates both promoter and enhancer elements, whereas TCF4 activates the enhancer in NE-4C cells but not in JEG-3 cells. In JEG-3 cells, TCF4 indirectly represses both regulatory elements and could offset activation by NFIB when co-expressed. Previous studies demonstrate that both NFIB and TCF4 can repress gene expression in other contexts (Chen et al., 2014; Piper et al., 2014), perhaps through their interaction with other transcriptional regulators in a context-dependent manner. We propose that the vast array of potential outcomes at different regulatory elements that ultimately converge to regulate gene expression serves as a means of fine-tuning gene expression and accounts for the differences in gene regulation observed in *Nfib* and *Tcf4* knockout mice despite their similar genome-wide binding. Moreover, we postulate that this contributes to the disparity in phenotypes observed as well as different severity of shared phenotypes between *NFIB* haploinsufficiency and Pitt-Hopkins syndromes.

We acknowledge two limitations of our study. Given the widespread expression of NFIB and TCF4 in different cell types, utilizing bulk sequencing methodologies may have masked cell type-specific regulatory events or led to statistically insignificant effect sizes in our datasets, leading to false negatives. Second, our approach for assigning regulatory elements to their target genes could similarly benefit from the integration of high-resolution epigenomic data collected using single-cell technologies. Nevertheless, our study demonstrates significant convergence between NFIB and TCF4 in regulating neuronal differentiation during neocortical development. They converge by largely binding to the same regulatory elements but are nonetheless capable of exerting different effects on gene regulation.

## Materials and Methods

### Mouse models and tissue collection

Animal protocols were approved by the University of Queensland Animal Ethics Committee in accordance with the Australian Code of Practice for the Care and Use of Animals for Scientific Purposes, or by the Institutional Animal Care and Use Committee of Washington University in St. Louis in accordance with guidelines from the National Institutes of Health. *Nfib* knockout mice (Nfib^tm1Rmg^) (Steele-Perkins et al., 2005) were housed at the Queensland Brain Institute animal facility and maintained on a C57BL/6J background. Timed-pregnant mice were obtained by placing male and female mice together overnight with the following day designated as E0.5. Timed-pregnant CD-1 mice used for CUT&RUN experiments and Duolink PLAs were obtained from Charles River Laboratories and housed at Washington University’s animal core facility.

Both male and female embryos were used for analyses. Embryos used for immunofluorescence co-staining were fixed by immersion (E14.5 or younger) or via transcardial perfusion using 4% (w/v) paraformaldehyde in phosphate-buffered saline (PBS). Tissue used for RNA isolation or CUT&RUN were microdissected from embryos in ice-cold PBS or Hank’s balanced salt solution (HBSS; Gibco) and either snap frozen for RNA isolation or immediately dissociated for CUT&RUN. Frozen tissue was stored at -80°C until RNA isolation.

### Immunofluorescence co-staining, image acquisition, and analysis

Post-fixed brains were embedded in 3% (w/v) BD Difco Noble agar and 50 µm coronal sections were cut using a vibratome (Leica Biosystems). Adjacent sections were mounted onto two separate slides and immunofluorescence co-staining was performed as previously described (Bunt et al., 2017). Briefly, antigen retrieval was performed by immersing mounted sections in citrate buffer (0.01 M sodium citrate, pH 6.0; 0.05% Tween-20) and heating to 110°C for four minutes. Immersed sections were gradually cooled to 95°C before removal from the citrate buffer. Sections were washed with PBS and blocked with 10% (v/v) normal donkey serum (Jackson ImmunoResearch) and 0.2% (v/v) Triton X-100 in PBS for one hour before overnight incubation at room temperature with primary antibodies diluted in 2% (v/v) normal donkey serum and 0.2% (v/v) Triton X-100 in PBS. Sections were subsequently washed with PBS and incubated with secondary antibodies diluted with 0.2% (v/v) Triton X-100 in PBS at room temperature for three hours. Following additional washes, the sections were counterstained with 4’,6-diamidino-2-phenylindole, dihydrochloride (DAPI; Invitrogen) and coverslipped using ProLong Gold antifade mountant (Invitrogen). The primary antibodies used were rabbit anti-NFIB (1:500; RRID: AB_1854424), mouse anti-TCF4 (1:500; RRID: AB_2864294), rat anti-CTIP2 (1:750; RRID: AB_2064130), and rat anti-SOX2 (1:2000; RRID: AB_3093719).

Specificity of the NFIB and TCF4 antibodies have been demonstrated using *Nfib* and *Tcf4* knockout mice, respectively (Chen et al., 2017; Wittmann et al., 2021). The secondary antibodies used were Alexa Fluor 488-conjugated donkey anti-mouse (1:500; RRID: AB_141607), Alexa Fluor 555-conjugated donkey anti-rabbit (1:500; RRID: AB_162543), and Alexa Fluor 647-conjugated donkey anti-rat (1:500; RRID: AB_2893138). Confocal images were acquired as z-stacks (5 images each spaced 1.33 μm apart) using a 20x objective on a Nikon AX-R confocal microscope using NIS-Elements Advanced Research software (v6.10.02). Maximum intensity projections were generated from each stack and pseudo-colored to permit overlay, and contrast-brightness enhanced for presentation using ImageJ (v2.14.0).

### RNA-seq and analysis

RNA isolation was performed using TRIzol reagent (Invitrogen) and subsequently treated with DNA-free DNA removal kit (Invitrogen). 1 μg of RNA was used to prepare mRNA libraries using the NEBNext Ultra II Directional mRNA-seq for Illumina kit and the NEBNext Poly(A) mRNA Magnetic Isolation Module (New England Biolabs). Libraries were pooled and sequenced (2×75 bp paired end) on an Illumina Hi-Seq 4000 at the University of Queensland genomics facility. Raw FASTQ files were trimmed using Trimmomatic (v0.39) and quantified using Salmon (v1.10.2; mm10). Differential expression analysis was performed in R (v4.4.0) using DESeq2 with default settings (v1.46.0) following removal of unwanted variation using RUVg via the RUVSeq package (v1.40.0) (Risso et al., 2014). Differentially expressed genes were defined as genes with *P*_adj_ ≤ 0.05 (*n* = 4 per genotype). *Tcf4* knockout RNA-seq data (GSE120463) (Mesman et al., 2020) was re-analyzed as described for *Nfib* knockout RNA-seq data. Gene ontology was performed using DAVID (v2021) with all genes from the associated DESeq2 analysis used as background. Gene ontology on shared gene lists was performed by restricting background genes to those that were included in both the *Nfib* and *Tcf4* DESeq2 analyses. Statistical significance was reported as the false discovery rate (FDR) determined after Fisher’s exact tests. ENCODE RNA-seq data analyzed in this study is listed in Table S13. Analysis was performed using DESeq2, normalized to counts of External RNA Controls Consortium (ERCC) spike-in reads.

### CUT&RUN and analysis

Activated papain (30 units/mL) (Worthington Biochemical Corporation) was prepared in HBSS according to the manufacturer’s instructions and subsequently used to digest freshly dissected tissue for 20 minutes at 37°C. Following this, tissue pieces were allowed to settle and the papain solution was replaced with ice-cold HBSS containing 0.2% (w/v) bovine serum albumin and 1% (v/v) fetal bovine serum. Gentle trituration was performed using fire-polished Pasteur pipets of descending diameters (firstly 600 μm, followed by 500 μm and 400 μm, each triturated at least five times) until complete dissociation of tissue. The supernatant containing dissociated cells was then filtered using a 70 μm Flowmi cell strainer (Bel-Art) and immediately used for CUT&RUN. CUT&RUN was performed using 500,000 cells per reaction and the CUTANA CUT&RUN kit (EpiCypher) according to the manufacturer’s instructions. 0.5 μg of the following antibodies were used for CUT&RUN: rabbit anti-NFIB (RRID: AB_1854424), rabbit anti-TCF4 (RRID: AB_2879076), and rabbit IgG negative control antibody (RRID: AB_2923178). We previously demonstrated that the NFIB antibody effectively immunoprecipitates NFIB (Bunt et al., 2017).

The NFIB antibody was also utilized previously for CUT&Tag (Zhang et al., 2026). The TCF4 antibody has been validated for ChIP-seq following CRISPR-mediated deletion of *Tcf4* (Terceros et al., 2026). We also verified specificity of the TCF4 antibody using TCF4 immunoprecipitation (Fig. S14). Isolated DNA (up to 5 ng) was used to make CUT&RUN libraries using the xGen DNA Library Prep Kit MC UNI and xGen UDI-UMI adapters (Integrated DNA Technologies). Libraries were pooled and sequenced (2×150 bp paired end) on an Illumina NovaSeq X Plus at the Genome Technology Access Center, Washington University McDonnell Genome Institute. Raw FASTQ files were trimmed using Trimmomatic (v0.36) and mapped to the mm10 genome using Bowtie2 (v2.3.5.1). PCR duplicates were removed using UMI-tools (v1.1.4) and peaks (*P* < 0.001) were called for individual samples using MACS (v3.0.0). Negative control rabbit IgG libraries were combined without removing PCR duplicates and used as control for peak calling. Irreproducible Discovery Rate (v2.0.4.2) was used to identify reproducible peaks between biological replicates (IDR < 0.05; *n* = 2). Motif analyses and peak annotation to identify the nearest gene were performed using HOMER (v5.1) (Heinz et al., 2010). Further processing of peaks was performed using bedtools (v2.31.1) and deepTools (v3.5.3). ENCODE ChIP-seq datasets used for peak annotation were obtained from the ENCODE portal (Kagda et al., 2025) and are listed in Table S13. ENCODE E14.5 forebrain ChromHMM chromatin states and predicted enhancer targets were obtained from the supplemental data of the original studies (Gorkin et al., 2020; Yu et al., 2025). We utilized ChromHMM chromatin states generated after pooling both replicates of the underlying ChIP-seq data. We performed permutation tests on expression-matched genes to determine whether NFIB and TCF4 binding to differentially expressed genes were significantly enriched. Briefly, 10,000 permutations were performed for each gene list, with wildtype controls from the E14.5 *Tcf4* knockout dataset used to identify expression-matched genes. To do this, all genes were filtered to retain only genes with a mean expression of at least 0.1 transcript per million (TPM), which we then divided into 20 equal-sized bins. For each permutation, randomly selected genes were obtained from the appropriate bins to match genes-of-interest, ensuring no overlap between permuted genes and genes-of-interest.

### Proximity ligation assays

Duolink PLAs (Sigma-Aldrich) were performed on 10 µm coronal sections obtained from E14.5 paraffin-embedded wildtype brains. Following deparaffinization and rehydration of tissue sections, heat-mediated antigen retrieval was performed in citrate buffer followed by tissue permeabilization in 0.2% (v/v) Triton X-100 in PBS for 30 minutes. PLA was subsequently performed according to the manufacturer’s instructions using the following primary antibodies: rabbit anti-NFIB (1:500; RRID: AB_1854424), mouse anti-TCF4 (1:500; RRID: AB_2864294), normal rabbit IgG (RRID: AB_1031062), and normal mouse IgG (RRID: AB_3675987). The normal rabbit and mouse IgG were used at an equimolar concentration to the NFIB and TCF4 antibodies, respectively. Confocal images were acquired as z-stacks (10 images each spaced 0.3 μm apart) using a 40x oil objective on a Nikon AX-R confocal microscope using NIS-Elements Advanced Research software (v6.10.02). Maximum intensity projections were generated from each stack and automatically quantified using ImageJ (v2.14.0). Statistical significance was determined using Student’s t-tests (*n* = 4). Representative images were pseudo-colored to permit overlay, and contrast-brightness enhanced for presentation in ImageJ.

### Cloning of plasmids for dual-luciferase reporter assays

The pPBCAG-*Tcf4*-IRES-GFP plasmid was cloned by excising *Tcf4* from pFUW-tetO-Tcf4 (Rosa et al., 2018) using EcoRI and inserting this into the EcoRI site of pPBCAG-IRES-GFP (Paolino et al., 2020). pFUW-tetO-Tcf4 was a gift from Filipe Pereira (Addgene plasmid #139806). The p.Arg580Trp *Tcf4* missense variant was subsequently generated through site-directed mutagenesis of pPBCAG-*Tcf4*-IRES-GFP using the Q5 site-directed mutagenesis kit (New England Biolabs) and the following primers: 5’-TGAGCGCCTGTGGGTCCGAGATA-3’ and 5’-CGGGCATTATTGGCCATCC-3’. Firefly luciferase reporter plasmids were cloned from C57BL6/J genomic DNA using the primers listed in Table S14. The forward and reverse primers include restrictions sites for NheI/SpeI and XhoI, respectively, which were used for cloning into pGL4.23 (Promega Corporation). For the *Nxph3* promoter, the primer pair was designed to amplify a region of 511 nucleotides immediately upstream of the *Nxph3* transcription start site. This corresponded to the promoter ATAC-seq peak (pooled samples data obtained from the supplemental data of Gorkin et al., 2020) that is bound by NFIB and TCF4 (Fig. 4C). The *Nxph3* distal enhancer corresponded to a region of 505 nucleotides downstream of *Slc35b1* encompassing an ATAC-seq peak that is bound by both NFIB and TCF4 (Fig. 4C). Plasmid sequences were verified by Sanger sequencing or Oxford Nanopore whole plasmid sequencing. Protein expression was validated by Western blot.

### Dual-luciferase reporter assays

NE-4C (RRID: CVCL_B063) and JEG-3 (RRID: CVCL_0363) cells were acquired from the American Type Culture Collection. Cell lines were not authenticated or tested for contamination during the study. NE-4C cells (10,000 cells per well) or JEG-3 cells (25,000 cells per well) were seeded into 96-well plates and maintained in Dulbecco’s Modified Eagle Medium (Gibco) supplemented with 10% (v/v) fetal bovine serum. Transfections were performed 24 hours after seeding. Briefly, each well was co-transfected with 60 ng of the firefly luciferase reporter plasmid and up to two of the following over-expression plasmids: negative control pCAG-IRES-GFP (40 ng when transfected alone or 20 ng when co-transfected with another over-expression plasmid), 20 ng of wildtype or mutant pCAG-*Nfib*-IRES-GFP (Schanze et al., 2018), or 20 ng of wildtype or mutant pPBCAG-*Tcf4*-IRES-GFP. 1 ng of the pNL1.1.TK plasmid (Promega Corporation) was co-transfected in all wells to normalize for transfection efficiency. Firefly and NanoLuc luciferase activity were measured 40 hours post-transfection using the Nano-Glo Dual-Luciferase Reporter Assay (Promega Corporation) and a BioTek Synergy HTX multimode plate reader (Agilent Technologies). To facilitate visualization and analysis, relative luciferase activity was multiplied by 1,000 and log_2_ transformed. Statistical significance was determined using two-way analysis of variance (ANOVA) to account for batch as a fixed effect. Post-hoc analyses were performed on the estimated marginal means using two-tailed t-tests and Holm-Bonferroni correction. *n* = 5 and 7 for each condition tested with the *Nxph3* promoter in JEG-3 cells and NE-4C cells, respectively. *n* = 4 and 6 for each condition tested with the *Nxph3* enhancer in JEG-3 cells and NE-4C cells, respectively.

## Acknowledgements

The authors thank Janette Edson at the University of Queensland for assistance with sequencing of the *Nfib* knockout RNA-seq libraries. Confocal images were acquired on a Nikon AX-R confocal microscope purchased with support from the Office of Research Infrastructure Programs, a part of the NIH Office of the Director under grant OD030233 at the Washington University Center for Cellular Imaging core facility. We also acknowledge the Research Infrastructure Services group at Washington University in St. Louis for providing computational resources and services.

## Data and resource availability

RNA-seq and CUT&RUN data generated in this study or re-analyzed will be publicly accessible upon manuscript acceptance. Plasmids generated in this study are available upon request.

## Author Contributions

J.W.C.L. and L.J.R. conceived and designed the study. J.W.C.L., E.H., N.A.M., J.E.A., S.P., and J.B. performed the experiments. J.W.C.L., E.H., N.A.M., and J.E.A. analyzed the data with H.B. providing support for software. J.B. and L.J.R. supervised the investigation. J.W.C.L. wrote the first draft of the manuscript and all authors edited and provided comments on the manuscript.

## Supplementary figures

**Figure S1.**
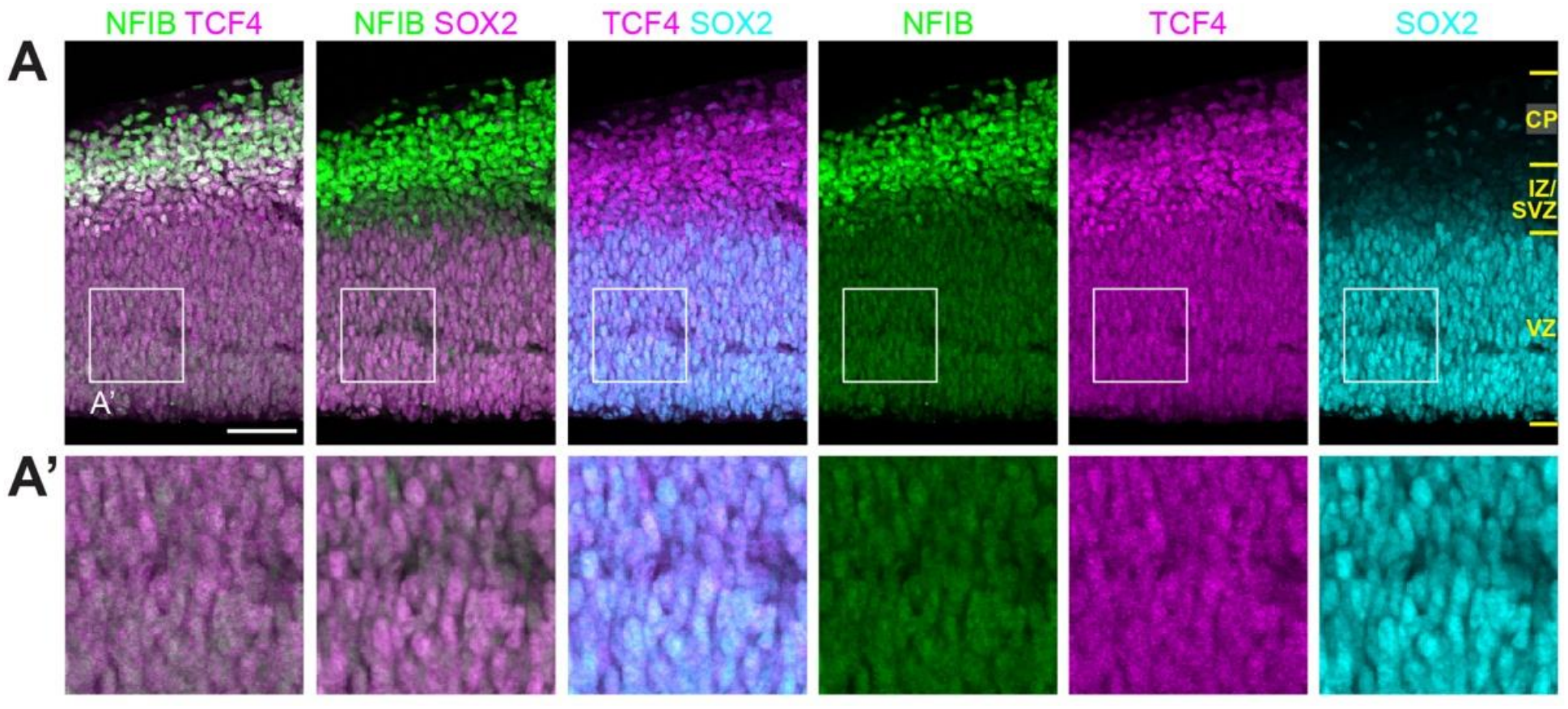
NFIB, TCF4, and SOX2 are co-expressed in the E13.5 neocortex. Adjacent coronal sections from those in Fig. 1 were stained for NFIB (green), TCF4 (magenta), and SOX2 (pseudo-colored magenta or cyan to enable visualization). Scale bar, 50 µm. (CP) Cortical plate, (IZ) intermediate zone, (SVZ) subventricular zone, (VZ) ventricular zone.

**Figure S2.**
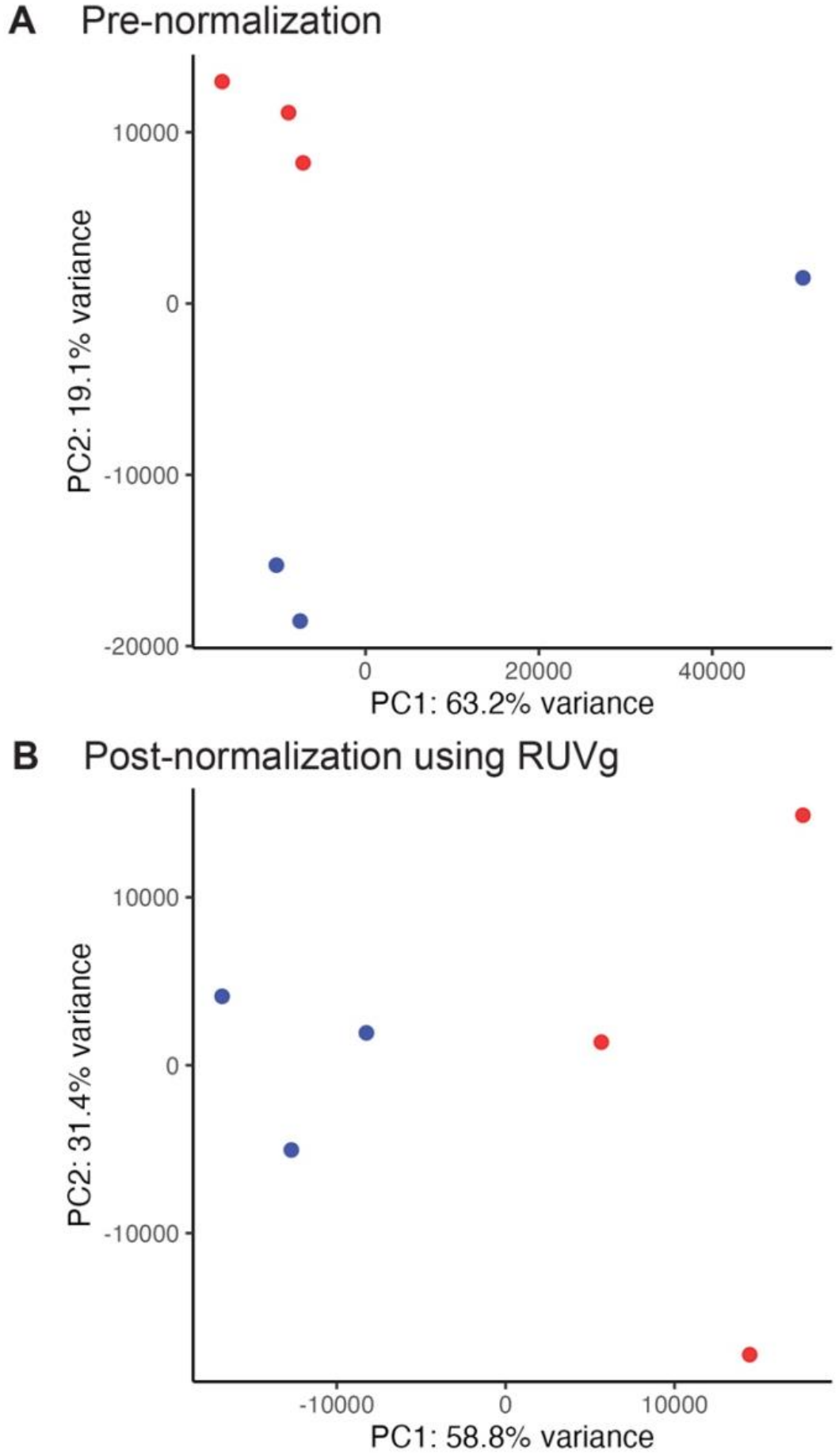
Principal component analysis plots of E14.5 *Tcf4* knockout samples (blue dots) and control littermates (red dots) before (*A*) and after (*B*) normalization using RUVg.

**Figure S3.**
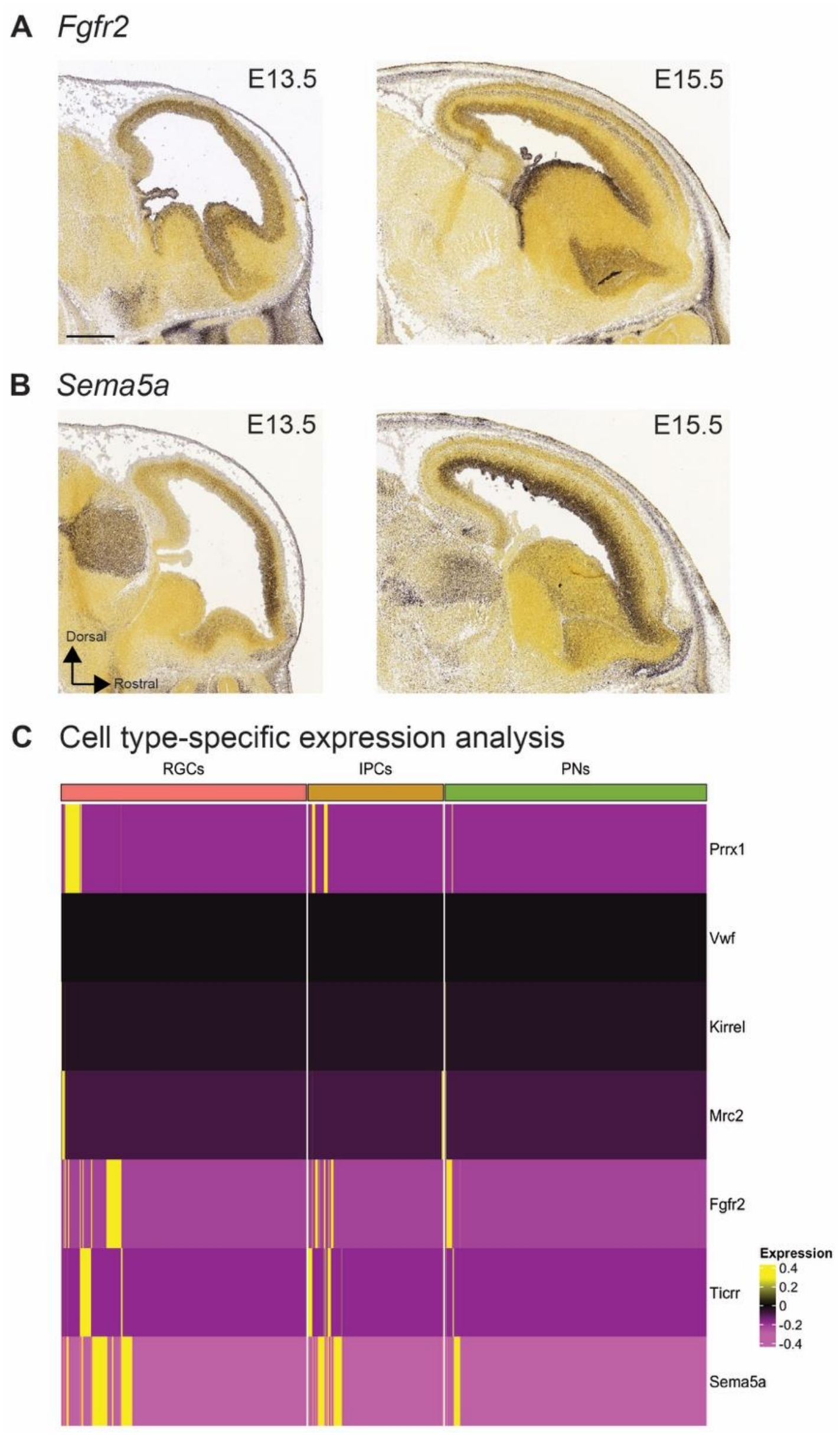
Shared upregulated genes between the E13.5 *Nfib* and E14.5 *Tcf4* knockout neocortices are expressed in radial glial cells and intermediate progenitor cells. (*A*,*B*) mRNA expression of *Fgfr2* (E13.5: developingmouse.brain-map.org/experiment/show/100049453; E15.5: developingmouse.brain-map.org/experiment/show/100085326) and *Sema5a* (E13.5: developingmouse.brain-map.org/experiment/show/100031956; E15.5: developingmouse.brain-map.org/experiment/show/113391531) in E13.5 and E15.5 sagittal sections. Images were downloaded from the Allen Developing Mouse Brain Atlas (Thompson et al., 2014). Scale bar, 500 µm. (*C*) Clustered heatmap depicting relative expression of the 7 shared upregulated genes in the three major cell types present within the E13.5 and E14.5 wildtype neocortex. Genes with variable expression were more prominently expressed in radial glial cells and intermediate progenitor cells as compared to postmitotic neurons. Gene expression data was obtained from a published single-cell RNA-seq (scRNA-seq) dataset (Di Bella et al., 2021). Z-score normalization was independently applied to each gene across all cells. Hierarchical clustering was then performed based on Euclidean distance with complete linkage. Outliers were capped to the minimum and maximum values indicated on the color scale. (RGCs) Radial glial cells, (IPCs) intermediate progenitor cells, (PNs) postmitotic neurons.

**Figure S4.**
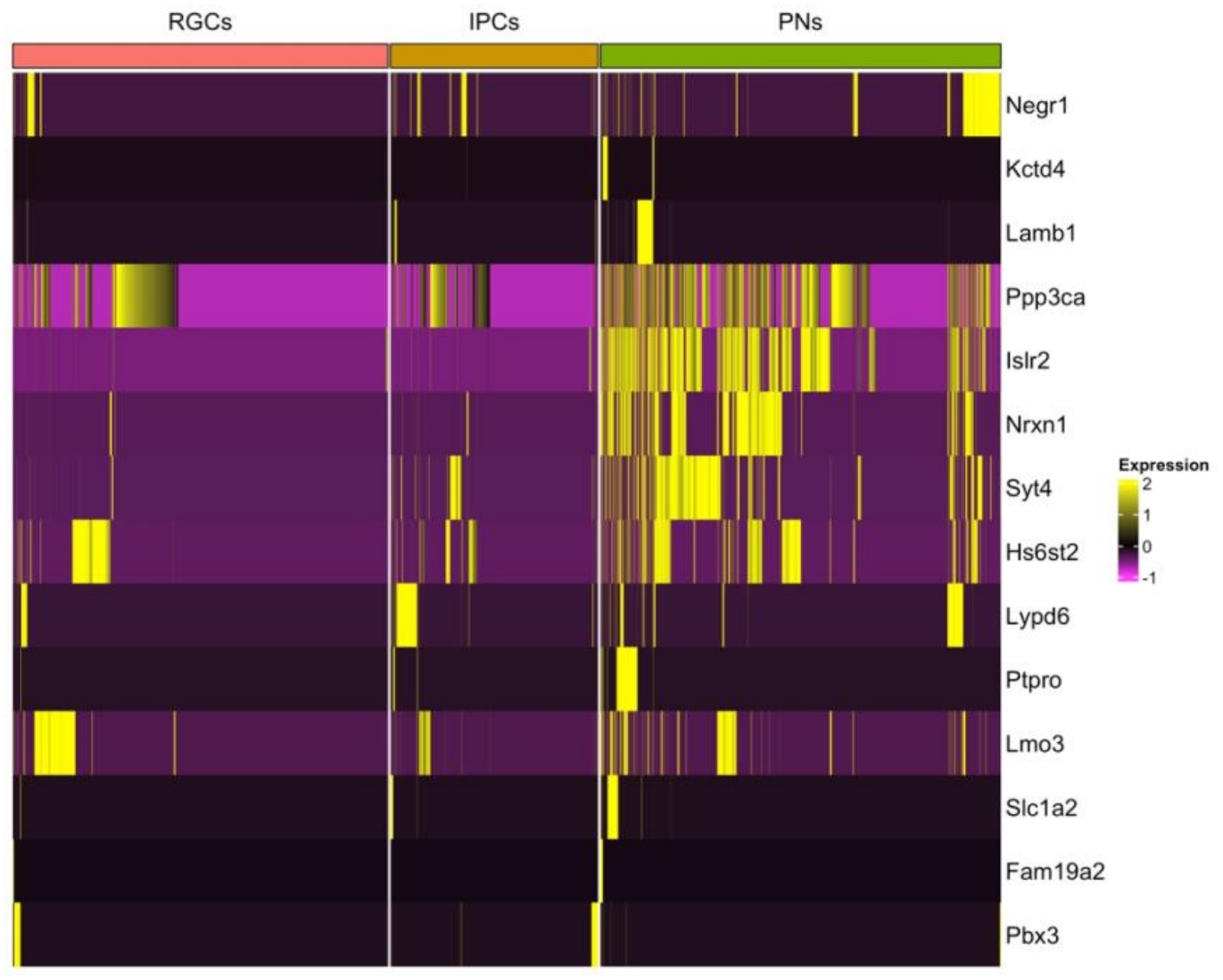
Clustered heatmap depicting relative expression of the 14 discordantly misregulated genes in the three major cell types present within the E13.5 and E14.5 wildtype neocortex. Many genes demonstrated higher expression in postmitotic neurons. Gene expression data was obtained from a published scRNA-seq dataset (Di Bella et al., 2021). Z-score normalization was independently applied to each gene across all cells. Hierarchical clustering was then performed based on Euclidean distance with complete linkage. Outliers were capped to the minimum and maximum values indicated on the color scale. (RGCs) Radial glial cells, (IPCs) intermediate progenitor cells, (PNs) postmitotic neurons.

**Figure S5.**
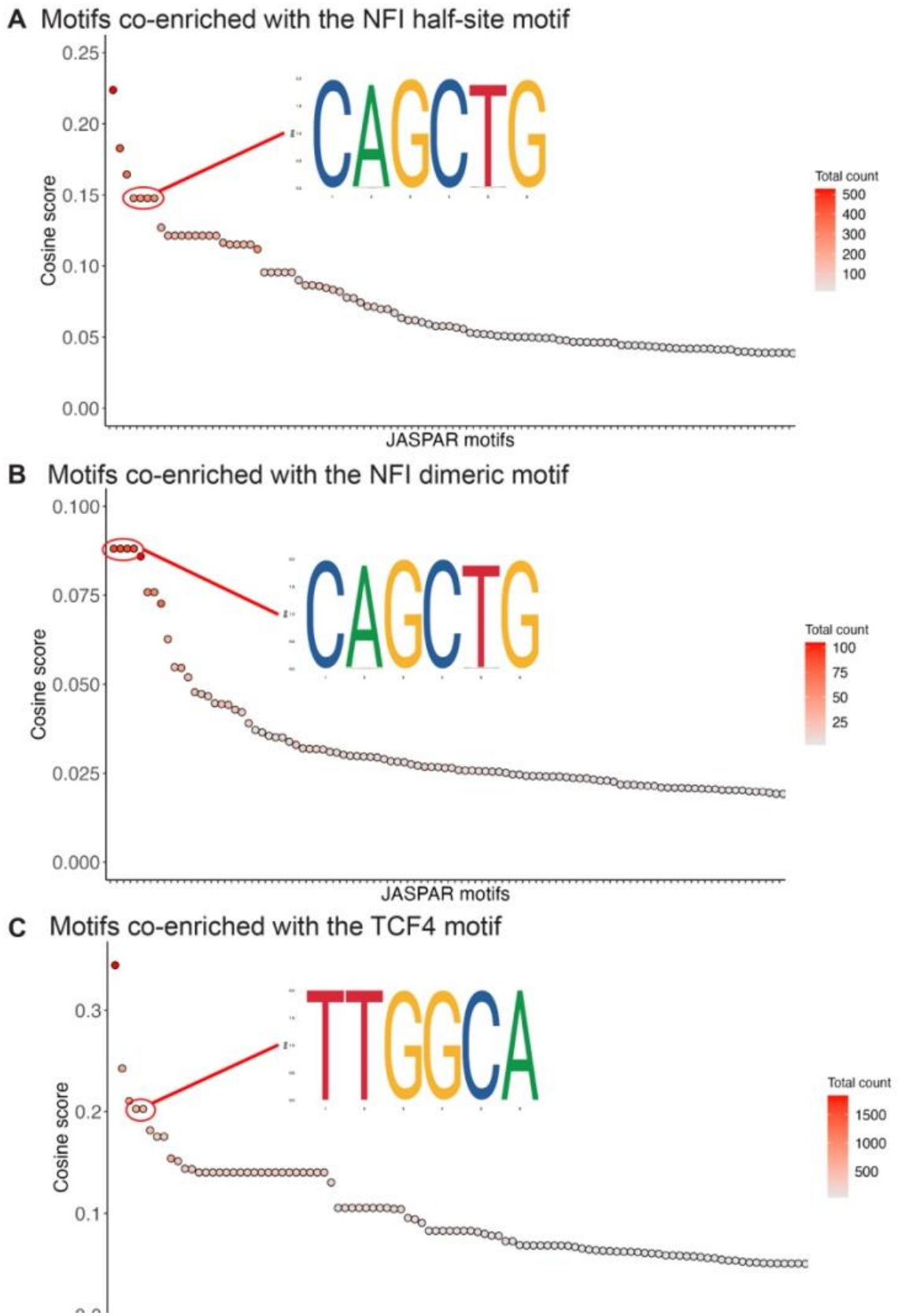
Market basket analyses performed using TF-COMB demonstrates co-occurrence of the NFI and E-box motifs at NFIB and TCF4 peaks. (*A*,*B*) Market basket analysis was performed on all NFIB peaks to examine co-occurrences of the NFI half-site (*A*) or dimeric motif (*B*) with all transcription factor motifs within the JASPAR 2024 CORE vertebrate database (Bentsen et al., 2022; Rauluseviciute et al., 2024). Enrichment of the E-box motif is highlighted. (*C*) Market basket analysis performed on all TCF4 peaks to examine motif co-occurrences with the TCF4 E-box motif. The enriched NFI half-site motif is highlighted.

**Figure S6.**
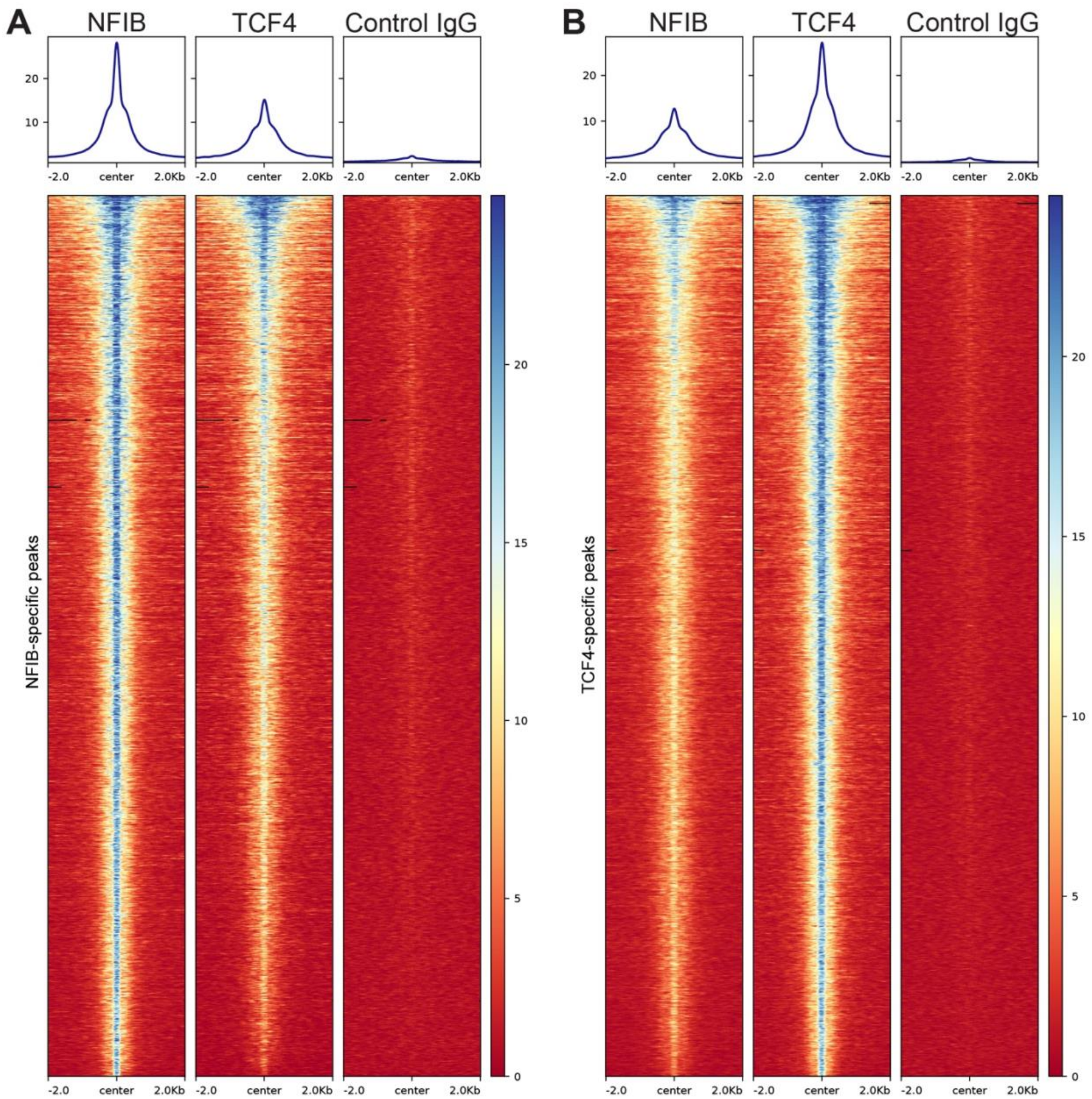
Weak TCF4 and NFIB binding is observed at NFIB-specific and TCF4-specific peaks, respectively. (*A*,*B*) Heatmaps depicting sequenced fragment density at NFIB-specific (*A*) and TCF4-specific (*B*) CUT&RUN peaks. Compared to IgG control, TCF4 fragments were enriched at NFIB peaks and vice versa but this weaker binding was insufficient to meet the cutoff used to identify reproducible peaks. Genomic regions were sorted by mean pileup of fragments across all samples. Black bars indicate genomic regions without any mapped fragments.

**Figure S7.**
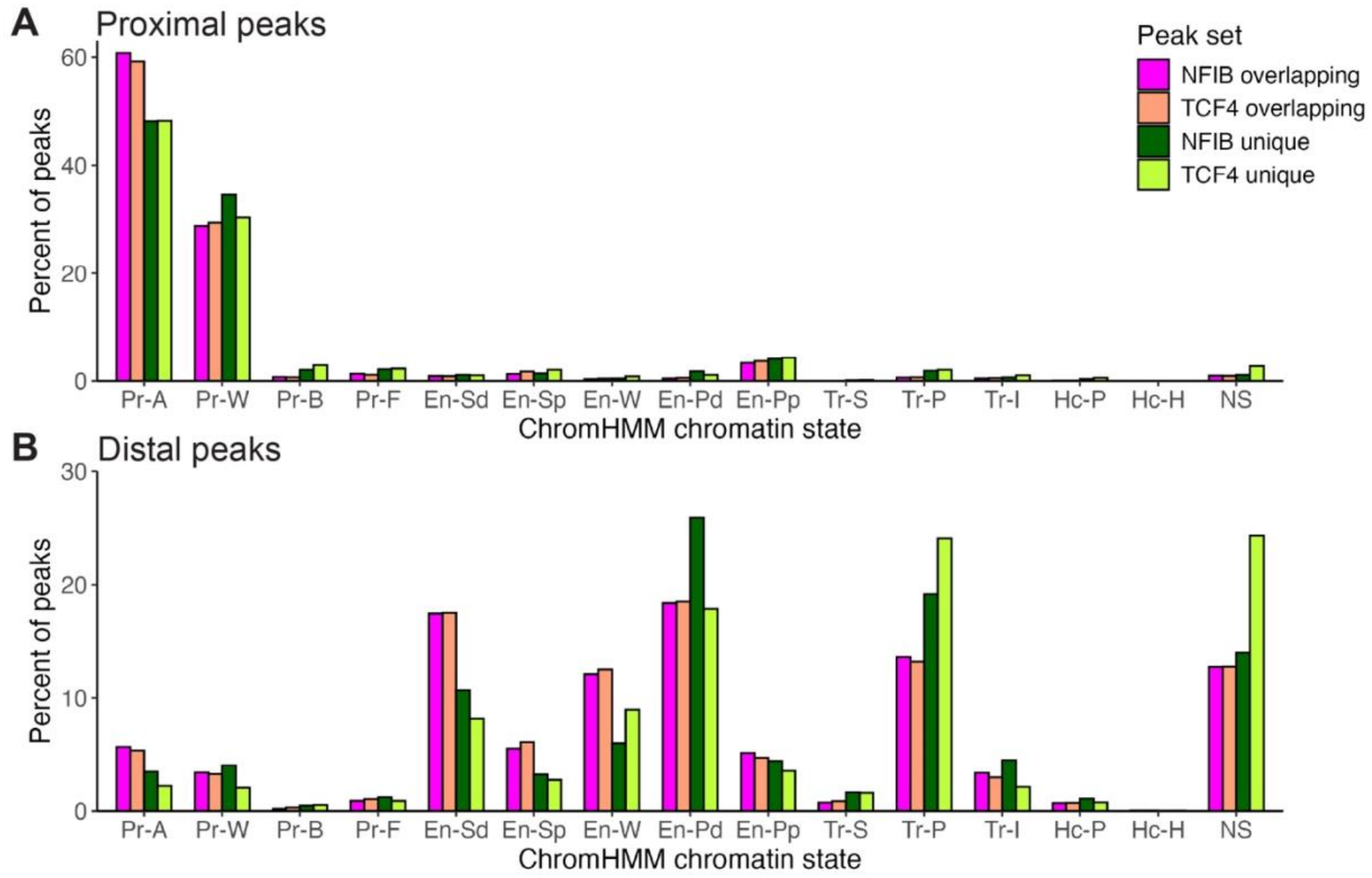
NFIB and TCF4 binding is associated with active promoters and distal enhancers. Overlapping and unique peaks were categorized as gene proximal (*A*) or distal (*B*) and subsequently assigned a ChromHMM chromatin state from ENCODE E14.5 forebrain data (Gorkin et al., 2020). (Pr-A) active promoter, (Pr-W) weak promoter, (Pr-B) bivalent promoter, (Pr-F) promoter flanking, (En-Sd) strong distal enhancer, (En-Sp) strong proximal enhancer, (En-W) weak distal enhancer, (En-Pd) poised distal enhancer, (En-Pp) poised proximal enhancer, (Tr-S) strong transcription, (Tr-P) permissive transcription, (Tr-I) transcription initiation, (Hc-P) Polycomb-associated heterochromatin, (Hc-H) H3K9me3-associated heterochromatin, (NS) no signal.

**Figure S8.**
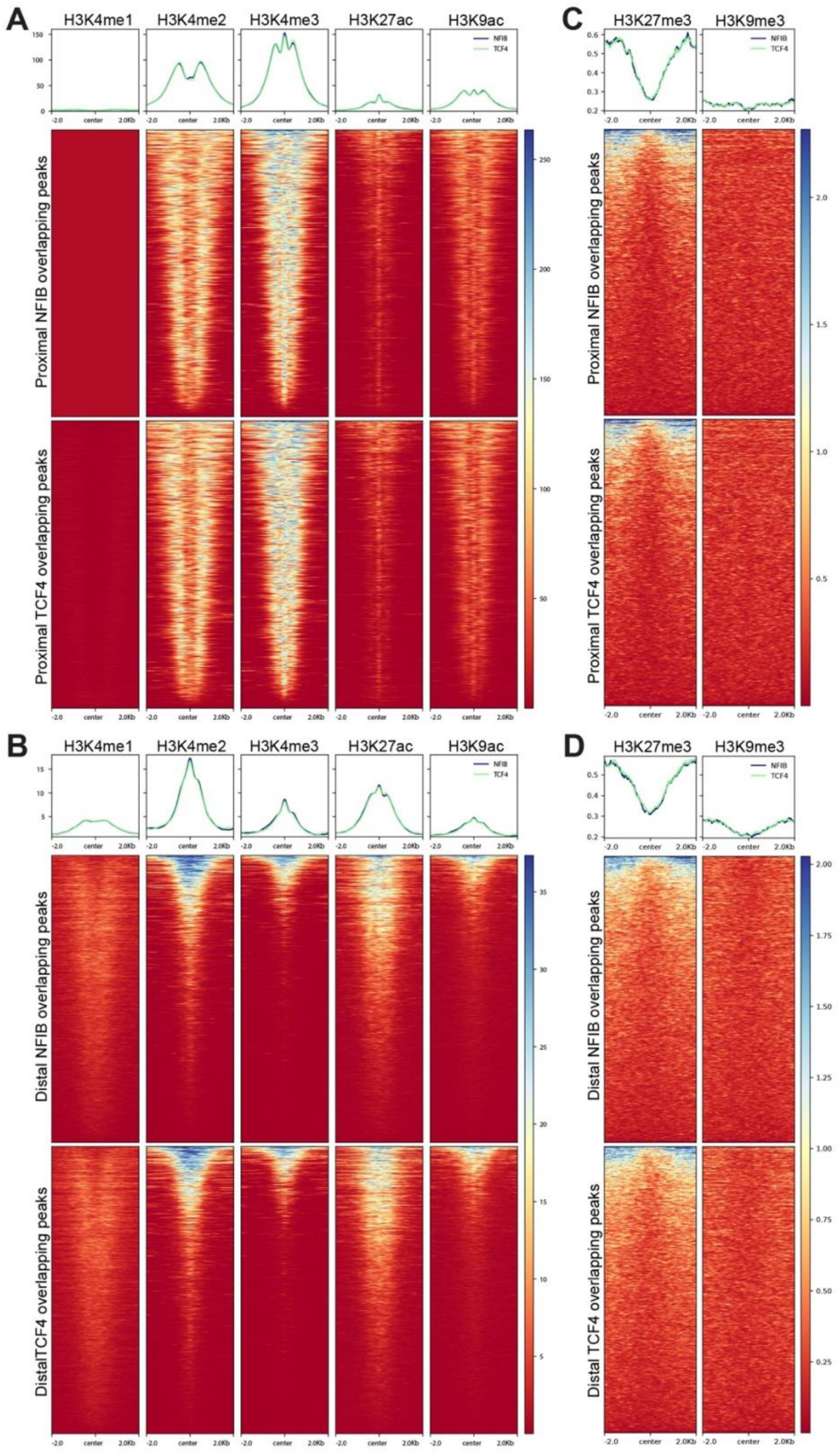
NFIB and TCF4 overlapping peaks are enriched for histone modifications associated with active promoters and distal enhancers. Profile plots and underlying heatmaps depict enrichment of histone modifications from ENCODE E14.5 forebrain ChIP-seq datasets. Genomic regions were sorted by mean pileup of fragments across all depicted histone modifications. (*A*,*B*) Enrichment of active histone modifications overlapping NFIB and TCF4 gene proximal (*A*) and distal (*B*) peaks. (*C*,*D*) Repressive histone modifications are not enriched at overlapping NFIB and TCF4 gene proximal (*C*) and distal (*D*) peaks.

**Figure S9.**
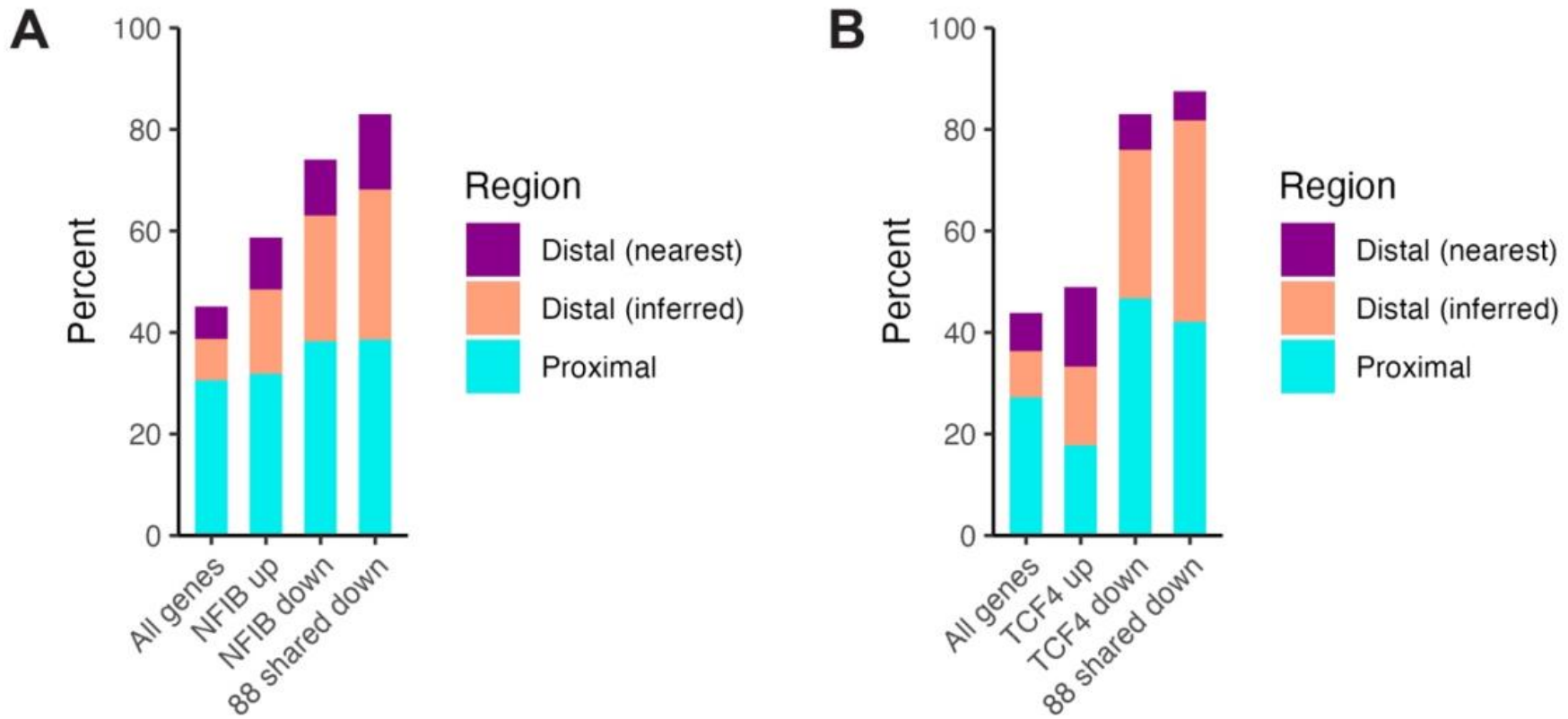
NFIB and TCF4 directly bind to most downregulated genes in the *Nfib* and *Tcf4* knockout neocortices, respectively. Bar charts depict the percentage of genes within each gene set that are directly bound by NFIB (*A*) or TCF4 (*B*) at gene proximal (aqua blue bars) or distal regulatory elements (salmon pink or dark purple bars). A gene is defined as proximally bound if proximal binding is detected regardless of whether distal binding was also observed. Distal bound genes are defined as genes with only distal binding and no proximal binding. Distal bound genes are grouped based on whether they were previously inferred (salmon pink bars) from the integration of various ENCODE datasets (Gorkin et al., 2020; Yu et al., 2025) or assigned to a gene using the nearest gene approach (dark purple bars).

**Figure S10.**
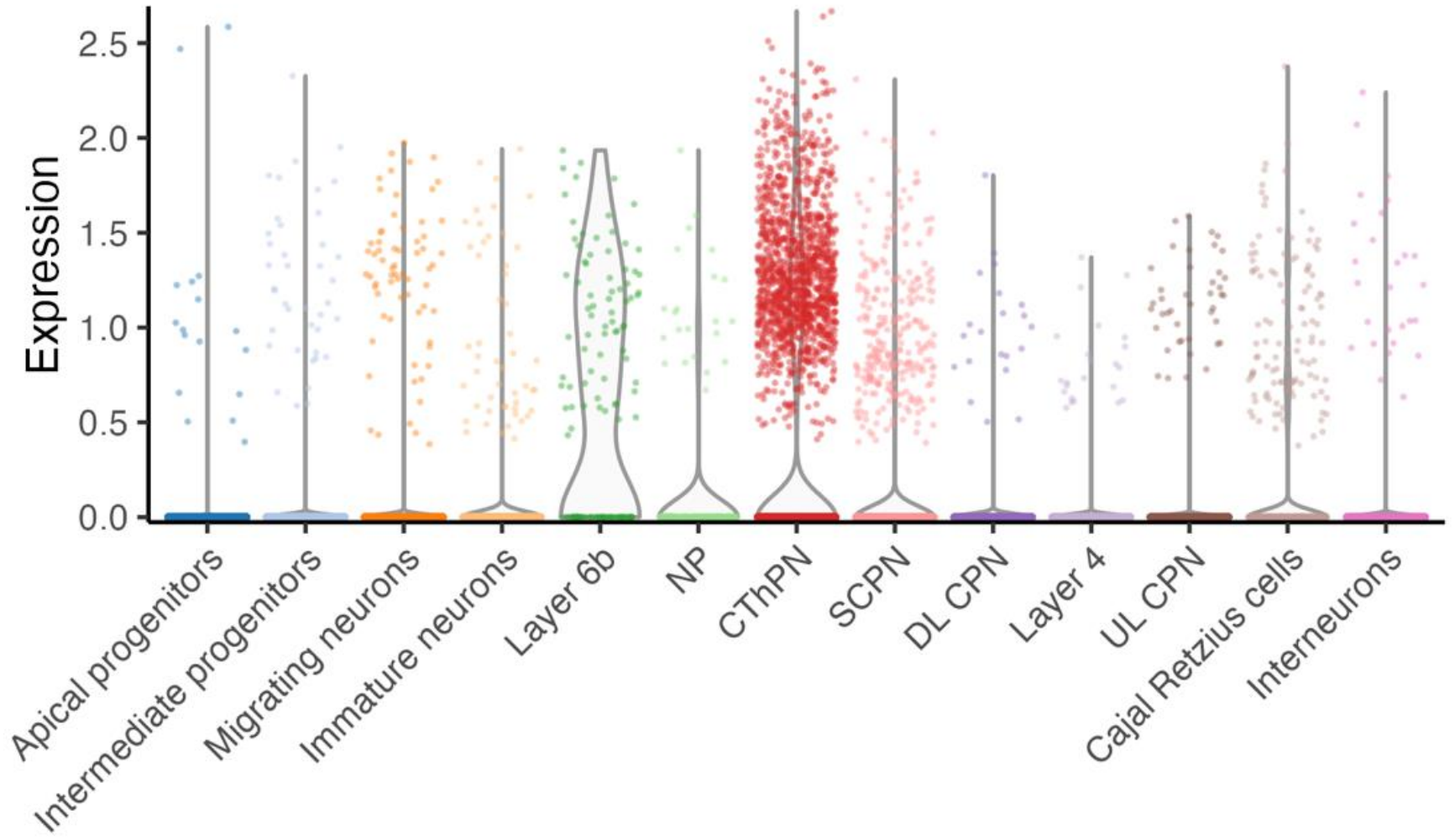
*Nxph3* is preferentially expressed in layer 6 (CThPN) and subplate (Layer 6b) cells during neocortical development. A published scRNA-seq dataset was filtered to keep only the depicted cell types throughout embryonic development (Di Bella et al., 2021). The violin plot depicts normalized expression of *Nxph3* across different cell types. (CThPN) layer 6 corticothalamic neurons, (DL CPN) layer 5 callosal neurons, (NP) layer 6 near-projecting neurons, (SCPN) layer 5 subcerebral projection neurons, (UL CPN) layer 2/3 callosal neurons.

**Figure S11.**
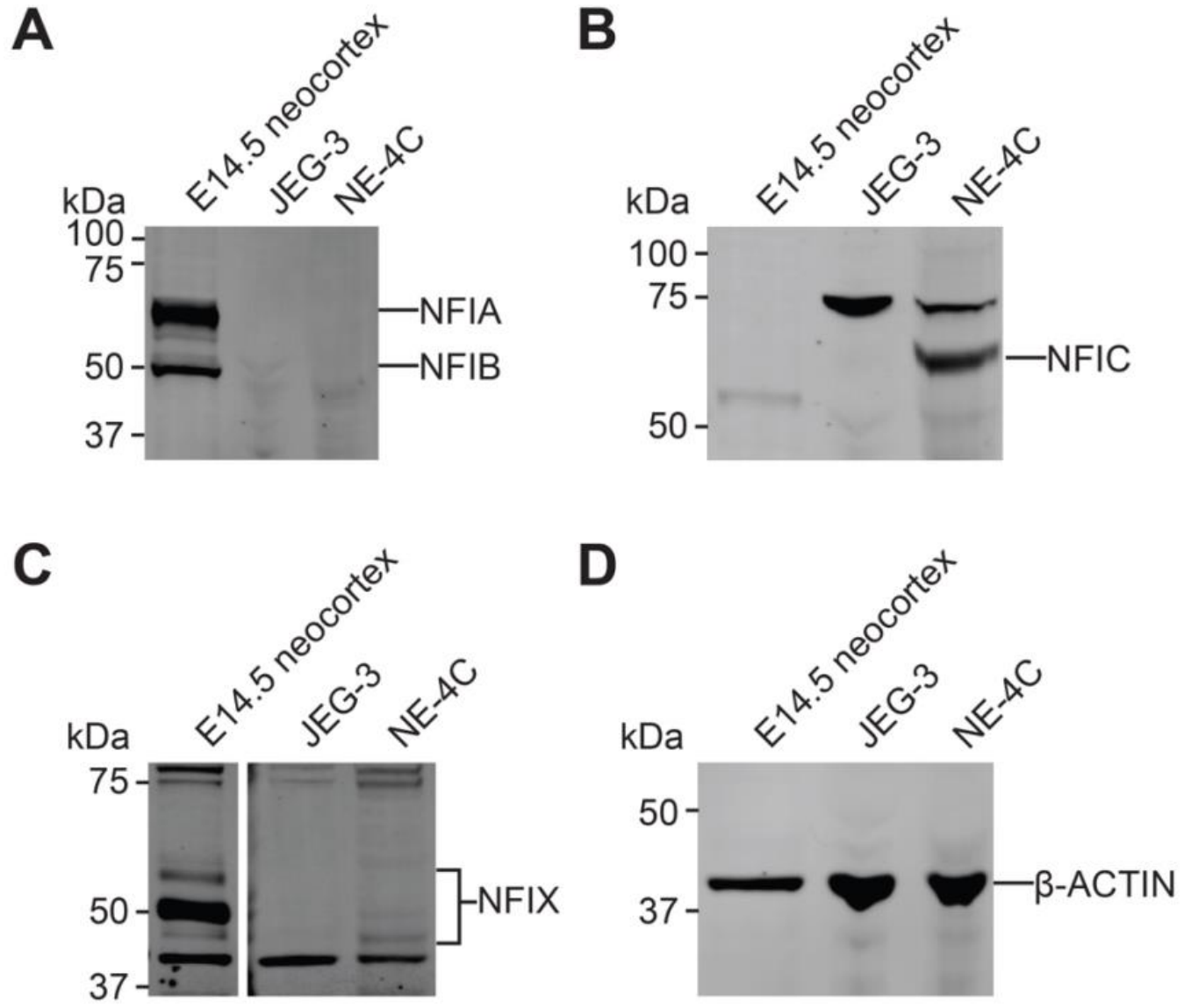
Representative Western blot images demonstrating endogenous expression of NFI transcription factors in whole cell lysates of JEG-3 and NE-4C cells. (*A*) NFIA and NFIB are lowly expressed in JEG-3 and NE-4C cells. E14.5 neocortical tissue lysate was used as a positive control. Immunoreactive bands for NFIA and NFIB in the positive control correspond to our previous observation with individual antibodies specific for NFIA or NFIB (Bunt et al., 2017). (B) NFIC expression is detected in NE-4C cells. An immunoreactive band of ∼60 kDa corresponding to NFIC (Huang et al., 2021; Rastogi et al., 2023; Sheng et al., 2021; Sumardika et al., 2019) was detected in NE-4C cells but not in JEG-3 cells. Although previous reports indicate that JEG-3 cells lowly express NFI transcription factors (Bruggemeier et al., 1990; Chaudhry et al., 1998), a second immunoreactive band of ∼75 kDa was detected in both cell lines, which may correspond to a post-translationally modified form of NFIC (Kane et al., 2002). (C) NFIX is lowly expressed in NE-4C cells. We observed an immunoreactive band of ∼55 kDa in E14.5 neocortical tissue lysate, which was previously reported with this antibody and validated by knockdown (Zhu et al., 2026). Additional immunoreactive bands of ∼50 kDa and ∼48 kDa were also observed with E14.5 neocortical tissue lysate and weakly immunoreactive in NE-4C cells. These bands correspond with the predicted molecular weight of mouse and human NFIX isoforms (UniProt IDs: P70257, Q14938). Additional bands of unknown identity were observed at ∼80, ∼75, and ∼42 kDa in all samples. (*D*) β-ACTIN was used as a loading control for Western blot. The primary antibodies used for Western blot were (*A*) mouse anti-pan-NFI recognizing NFIA and NFIB (1:200; RRID: AB_2618885), (*B*) rabbit anti-NFIC (1:1,000; RRID: AB_2681889), (*C*) mouse anti-NFIX (1:1,000; RRID: AB_2918732), and (*D*) mouse anti-β-ACTIN (1:1,000; RRID: AB_2273656). The secondary antibodies used were IRDye 680RD donkey anti-rabbit IgG (1:20,000; RRID: AB_10954442) and IRDye 800CW donkey anti-mouse IgG (1:20,000; RRID: AB_621847). Western blot images were acquired on an Odyssey CLx imager (LI-COR Biotech) using Image Studio (version 6.0) and brightness-contrast adjusted for visualization.

**Figure S12.**
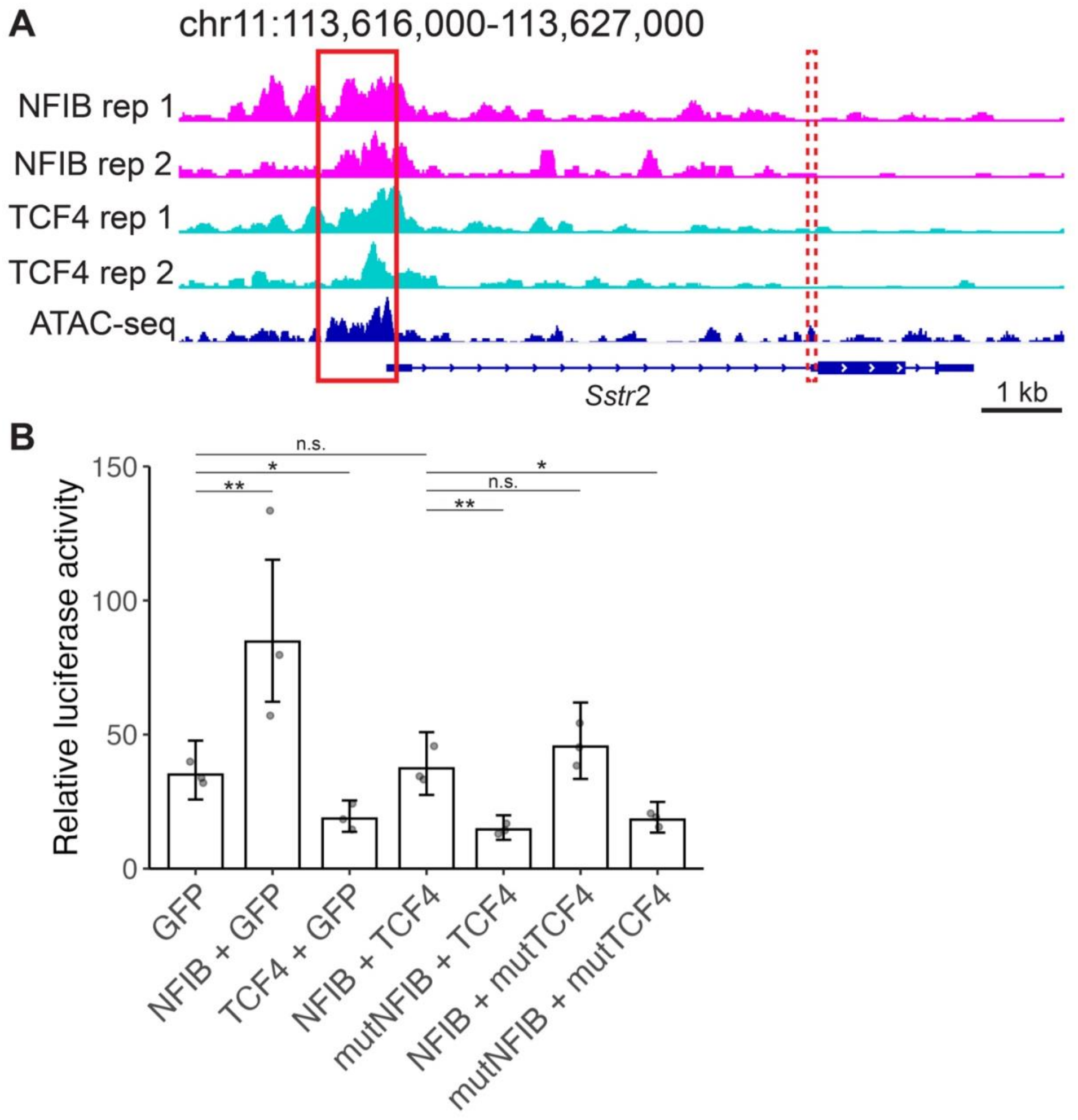
NFIB activates the *Sstr2* promoter in JEG-3 cells whereas TCF4 represses its activity. (*A*) Genome browser plot demonstrating NFIB and TCF4 binding at the *Sstr2* promoter. The ATAC-seq track represents ENCODE E14.5 forebrain ATAC-seq data (Gorkin et al., 2020). The solid red box (left) depicts the region cloned for luciferase assays. The dashed red box (right) depicts a potential regulatory element overlapping the exon-intron boundary that is just upstream of the protein coding sequence and is activated by TCF4 (Dorflinger et al., 1999). This region was previously thought to be the *Sstr2* promoter. (*B*) Relative luciferase activity of the *Sstr2* promoter in JEG-3 cells. Data represents mean ± 95% c.i. (*n* = 3 biological replicates). Two-tailed Student’s t-test in (*B*): (*) *P* = 0.025 for GFP versus TCF4 + GFP, 0.0151 for NFIB + TCF4 versus mutNFIB + mutTCF4, (**) *P* = 0.0043 for GFP versus NFIB + GFP, 0.0031 for NFIB + TCF4 versus mutNFIB + TCF4, (n.s.) *P* = 0.7546 for GFP versus NFIB + TCF4, 0.69 for NFIB + TCF4 versus NFIB + mutTCF4.

**Figure S13.**
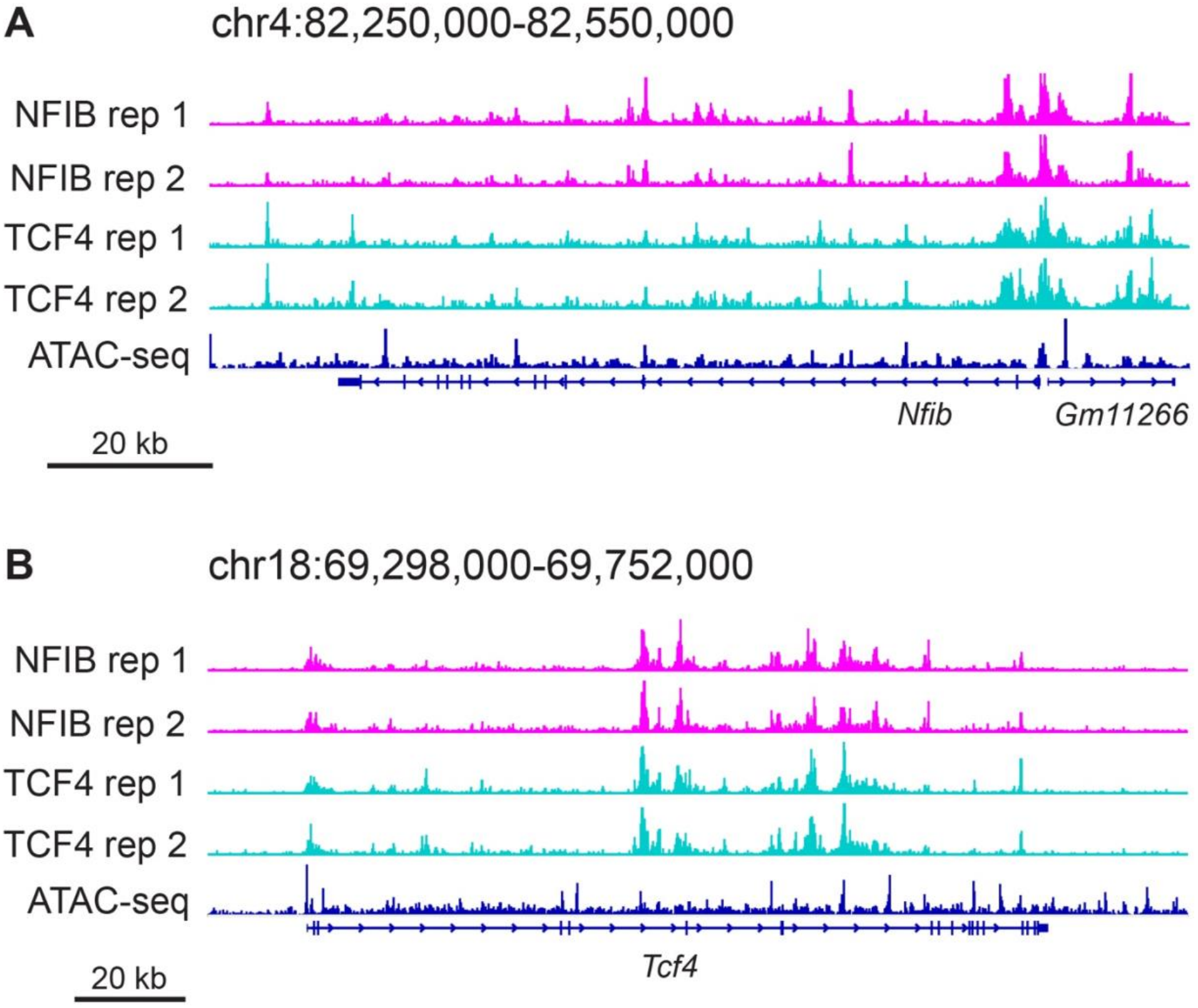
NFIB and TCF4 directly co-regulate each other’s expression. Genome browser plots depicting NFIB and TCF4 binding at the *Nfib* (*A*) and *Tcf4* (*B*) loci. The ATAC-seq track represents ENCODE E14.5 forebrain ATAC-seq data (Gorkin et al., 2020).

**Figure S14.**
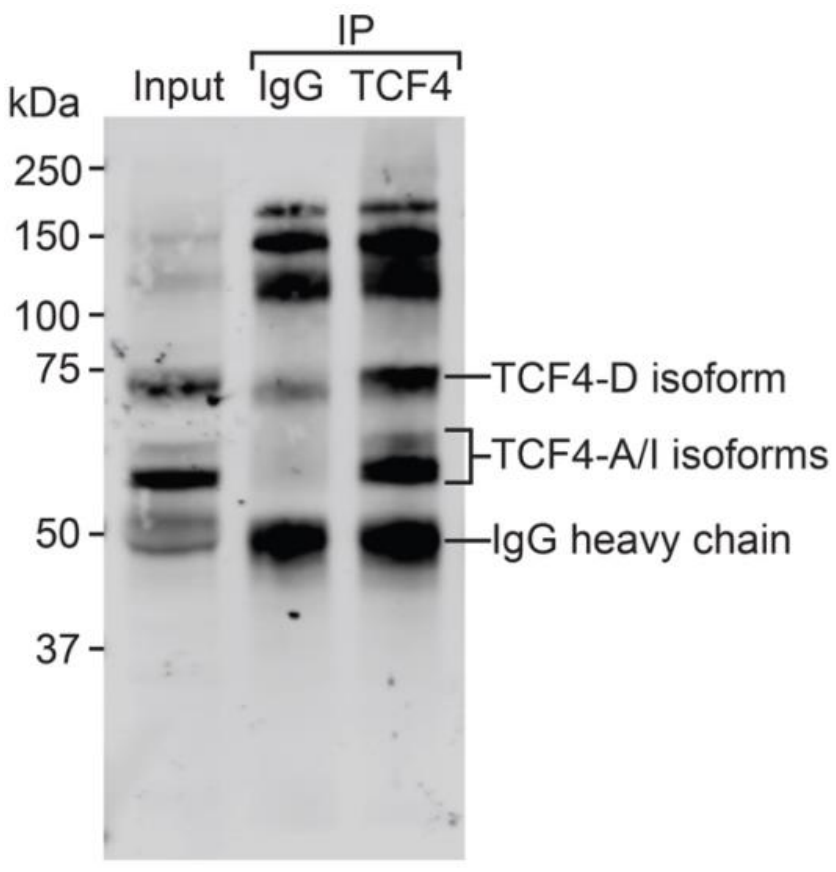
Representative Western blot image demonstrating specificity of the TCF4 antibody used for CUT&RUN. Immunoprecipitation of E14.5 neocortical tissue lysate was performed using 1 μg of rabbit anti-TCF4 antibody (RRID: AB_2879076) or control rabbit IgG (RRID: AB_2923178) and Dynabeads Protein A (Invitrogen). Antibodies used for Western blot were mouse anti-TCF4 (1:1,000; RRID: AB_2864294) and IRDye 800CW donkey anti-mouse IgG (1:20,000; RRID: AB_621847). Immunoreactive bands corresponding to the short TCF4-A/I and medium TCF4-D isoforms were observed (Sirp et al., 2022).

## Supplementary tables

**Table S1. Top 100 enriched Gene Ontology Biological Process terms associated with upregulated genes in E13.5 *Nfib* knockout mice.**

**Table S2. Top 100 enriched Gene Ontology Biological Process terms associated with downregulated genes in E13.5 *Nfib* knockout mice.**

**Table S3. Enriched Gene Ontology Biological Process terms associated with upregulated genes in E13.5 *Tcf4* knockout mice.**

**Table S4. Top 100 enriched Gene Ontology Biological Process terms associated with downregulated genes in E13.5 *Tcf4* knockout mice.**

**Table S5. DESeq2 results for shared upregulated genes in E13.5 *Nfib* knockout and E14.5 *Tcf4* knockout mice.**

**Table S6. DESeq2 results for shared downregulated genes in E13.5 *Nfib* knockout and E14.5 *Tcf4* knockout mice.**

**Table S7. Wilcoxon rank sum tests comparing expression of the 88 shared downregulated genes between major cell types in the neocortex at E13.5 and E14.5, obtained from a temporal single-cell atlas (Di Bella et al., 2021).** Differential expression testing was performed using Seurat (version 5.2.1).

**Table S8. Top 100 enriched Gene Ontology Biological Process terms associated with shared downregulated genes in E13.5 *Nfib* knockout mice and E14.5 *Tcf4* knockout mice.**

**Table S9. DESeq2 results of ENCODE RNA-seq data comparing E13.5 and E14.5 forebrain tissue, filtered to retain only the 88 shared downregulated genes.**

**Table S10. DESeq2 results for discordantly misregulated genes between E13.5 *Nfib* knockout and E14.5 *Tcf4* knockout mice.**

**Table S11. Annotation of NFIB CUT&RUN peaks.** For peaks assigned to multiple genes, the assigned genes (Ensembl_ID) and assignment method (Method) are listed in the same order.

**Table S12. Annotation of TCF4 CUT&RUN peaks.** For peaks assigned to multiple genes, the assigned genes (Ensembl_ID) and assignment method (Method) are listed in the same order.

**Table S13. ENCODE accession IDs of RNA-seq and ChIP-seq datasets used in analyses.**

**Table S14. Primer sequences used to clone luciferase reporter plasmids.**

